# Capturing volumetric dynamics at high speed in the brain by confocal light field microscopy

**DOI:** 10.1101/2020.01.04.890624

**Authors:** Zhenkun Zhang, Lu Bai, Lin Cong, Peng Yu, Tianlei Zhang, Wanzhuo Shi, Funing Li, Jiulin Du, Kai Wang

**Author notes:** Correspondence should be addressed to K. W. These authors contributed equally to this work (Zhenkun Zhang, Lu Bai, Lin Cong).

## Abstract

Neural network performs complex computations through coordinating collective neural dynamics that are fast and in three-dimensions. Meanwhile, its proper function relies on its 3D supporting environment, including the highly dynamic vascular system that drives energy and material flow. Better understanding of these processes requires methods to capture fast volumetric dynamics in thick tissue. This becomes challenging due to the trade-off between speed and optical sectioning capability in conventional imaging techniques. Here we present a new imaging method, confocal light field microscopy, to enable fast volumetric imaging deep into brain. We demonstrated the power of this method by recording whole brain calcium transients in freely swimming larval zebrafish and observed behaviorally correlated activities on single neurons during its prey capture. Furthermore, we captured neural activities and circulating blood cells over a volume ⌀ 800 μm × 150 μm at 70 Hz and up to 600 μm deep in the mice brain.

## Introduction

Brain develops complex functions by recruiting large ensembles of cells with diverse functions and coordinating them precisely in time. Better understanding of these processes requires tools to capture the dynamic organization of the involved cells at biologically relevant spatial and temporal resolution, and in their native environment. Traditional tools developed for in-vivo imaging in the brain, such as confocal microscopy and two-photon scanning microscopy^1^, are based on point scanning scheme and too slow to study fast volumetric dynamics. Therefore, a number of faster imaging strategies have been introduced to meet this challenge, including new scanning mechanisms^2–4^, spatial and temporal multiplexing^5–9^ and exploiting the sparsity of samples^2, 4, 10–12^. However, the fluorescence saturation and animals’ tolerance of laser power still significantly limit the achievable imaging throughput^13–16^. Furthermore, capturing extremely fast dynamics, such as circulating blood cells, over an extended 3D volume has not yet been achieved. Light sheet microscope relaxes this trade-off by increasing parallelization in image collection and demonstrated high speed imaging in live cells^17^ and small animals, such as C. elegans^18, 19^ and larval zebrafish^20, 21^, but it is still challenging to get optimal performance when imaging deep into scattering mammalian brain^22^.

Light field microscopy (LFM) completely parallelizes the fluorescence collection and could capture information over an entire volume simultaneously in a single camera frame. Therefore, it is potentially the most favorable scheme for high speed three-dimensional (3D) imaging of fast dynamics in large biological tissues^23–26^. However, conventional LFMs lack optical sectioning capability and could not provide optimal spatial resolution and signal to noise ratio (SNR) when imaging into large brains. Special algorithms have been developed to extract neural signals in scattering mouse brain imaged by LFM^27–29^, but they are still very susceptible to strong background-induced Poisson noise and also not generally applicable to capturing dynamics involving rapid structural changes. Selective excitation over a small volume from the side to get higher resolution and reduce reconstruction artifacts in LFM has also been demonstrated^30^, but this optical configuration cannot be applied to mammalian brain, or to image freely moving animals.

Here, we presented a new type of LFM, termed Confocal LFM, to enable background-free fast volumetric imaging deep in the brain. Confocal LFM is equipped with a novel generalized form of confocal detection scheme to ensure selective and efficient signal collection from the in-focus volume, which is very different from conventional confocal detection designs that only extract signal from one single imaging plane at a time. The elimination of background, also known as the optical sectioning capability, preserves high signal to noise ratio and reduces reconstruction artifacts. To demonstrate the power of this method, we reported results from several example applications. We achieved blur-free imaging of the whole brain of a freely behaving larval zebrafish at 2 × 2 × 2.5 μm^3^ spatial resolution and over Ф 800 μm × 200 μm imaging volume. We also demonstrated calcium imaging of populations of neurons at greatly improved signal to background ratio and capturing circulating blood cells at depths up to 600 μm and at speed of 70 Hz in awake mouse brain. The introduction of optical sectioning capability to LFM significantly extends its application in studying various rapid volumetric dynamics deep in the brain.

## Results

### Confocal Light Field Microscopy

Confocal LFM uses the same optical design in our previously reported eXtended field of view LFM (XLFM)^31^, which was closely related to Shack-Hartmann wavefront sensor in adaptive optical microscope^32^ and placed micro-lens array at the objective’s conjugate pupil plane^33, 34^. This configuration eliminates reconstruction artifacts near the focal plane in traditional LFM^30, 31, 34^. Different from conventional LFM that employs wide-field illumination, in Confocal LFM we shaped the excitation laser beam into a plane, which goes through a thin slit in the center of a specially designed mask and a micro-lens conjugated to the center of the objective’s back pupil and selectively illuminate an axial sample plane (x-z plane) under the objective (Figure 1a). Fluorescence image formed by this central micro-lens was spatially filtered by the same slit on the mask and then conjugated to the imaging camera (Figure 1a, Supplementary Video 1, Supplementary Figure 1). The excitation light sheet, together with the mask, were scanned in y direction at the same speed by coordinating the galvo mirror and the rotating stage to cover a volume (Supplementary Figure 2). In this way, the scanning excitation light sheet, scanning slit on the mask, central micro-lens and the objective worked together as a conventional scanning line confocal microscope. Meanwhile, images formed by other micro-lenses were also spatially filtered by apertures on the same mask but of different shapes depending on the location of their corresponding micro-lenses (Figure 1a, Supplementary Figure 3). They were then conjugated to the imaging camera simultaneously and integrated during the scanning. At the end of each scan, camera read out one raw image for the volume reconstruction later.

**Figure 1.**
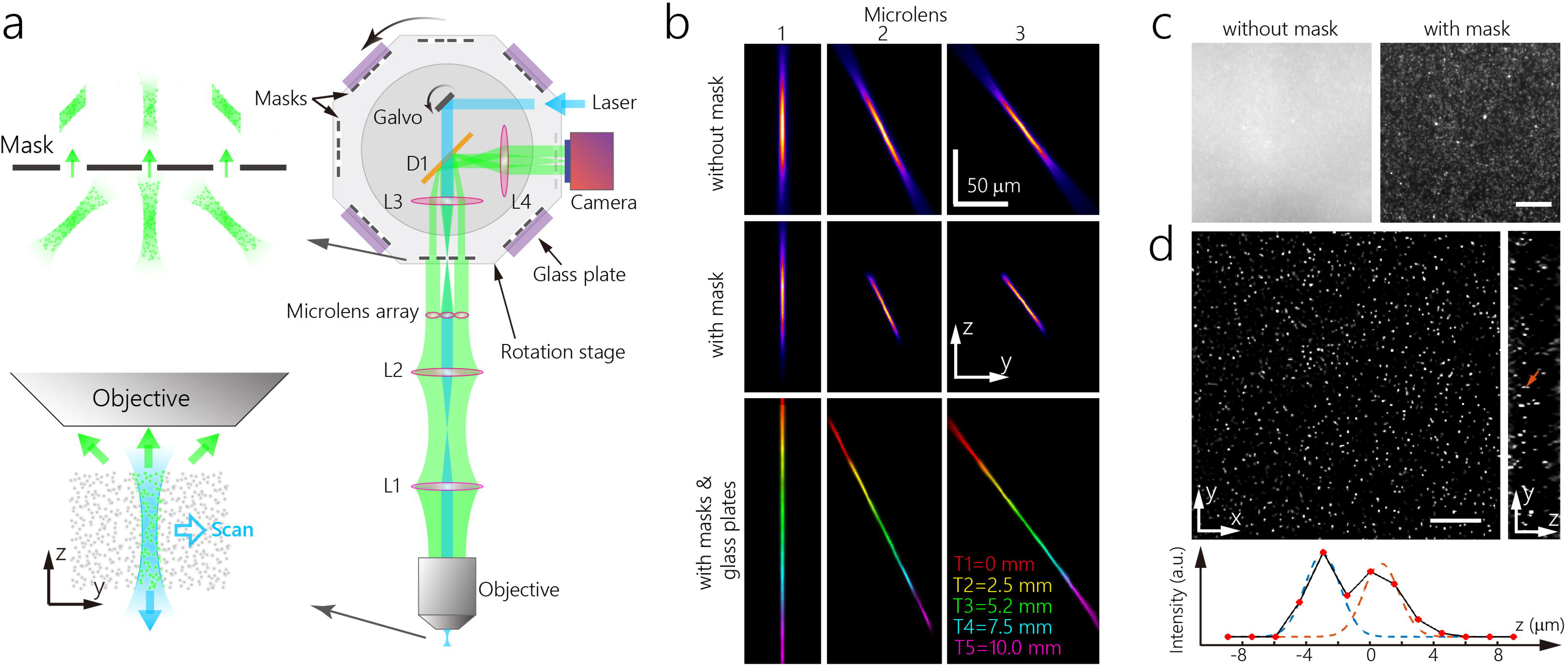
Design and characterization of Confocal LFM. (a) Overview (right) and zoom-in views (left) of the optical system. L1-L4: achromatic optical relay lenses; DI: short pass dichroic mirror. Zoom in views are vertical scanning light sheet in the sample plane (bottom) and its images formed by micro-lens array being spatially filtered by an optical mask (top), (b) Projection views of experimentally measured PSFs of three example micro-lenses with/without mask and glass plates (Supplementary Figure 4,5). Glass plates of different thickness (0,2.5 mm, 5.2 mm, 7.6 mm and 10 mm) shifted their PSFs (red, yellow, green, cyan and magenta) to different z locations to cover a larger axial range continuously, (c) Zoom-in views of raw images collected in LFM with/without mask when imaging fluorescent beads embedded in 3D agarose (Supplementary Figure 10). (d) representing images of fluorescent beads from single planes in 3D reconstructed volume. The yellow arrow indicates two fluorescent beads that are closely spaced in axial direction. Their pixel profile in z direction is plotted at the bottom and fitted by two Gaussian functions, indicating an estimated inter-beads distance of 3.4 *μ* m.

This confocal detection scheme is highly efficient as it does not discard any fluorescence signal from in-focus volume but blocks nearly all out-of-focus background. It can also run at high speed. The camera’s frame rate, rather than the scanning of excitation laser beam and mask, limits the volumetric imaging speed.

To further extend the capability of Confocal LFM to offer optimal spatial resolution over larger axial range, we developed a fast remote focusing mechanism to rapidly switch the imaging volume between multiple discrete axial locations. This was achieved by inserting one out of a series of glass plates of different thicknesses between the mask and the micro-lens array (Figure 1a, Supplementary Video 1, Supplementary Figure 1). Glass plates effectively changed the length of optical path and therefore shifted the conjugate plane of the mask to different axial locations under the objective. In fact, we could carefully choose parameters of these glass plates to continuously cover a larger volume by stitching together multiple shallower regions captured in each scan.

We characterized Confocal LFM by imaging and reconstructing fluorescent beads embedded in agarose gel. In one Confocal LFM implementation, the system has an effective Numerical Aperture (NA) of 0.13 for each micro-lens and can achieve lateral spatial resolution of 2 μm in 45 μm axial range (Figure 1b, Supplementary Figure 4). Imaging volumes could be raster scanned at five different axial locations to cover a ⌀ 800 μm × 200 μm total volume (Figure 1b, Supplementary Figure 5). We characterized the lateral resolution limit by measuring the Full Width at Half Maximum (FWHM) of the PSF at the in-focus plane (Supplementary Figure 6) which was about 2.1 μm. The axial resolution limit was determined by the lateral size and tilt angle of the PSFs measured through the outermost micro-lenses, which was about 2.5 μm (Supplementary Figure 6). By correcting system aberration, we could achieve this diffraction limited performance in reconstruction over the entire imaging volume (Supplementary Figure 7-9). We further confirmed that our system could at least clearly resolve two fluorescent beads separated by 3.4 μm in axial direction (Figure 1c). All above mentioned imaging performance could be preserved in Confocal LFM, but not in conventional LFM, due to the overwhelming background noise when imaging optically thick samples (Figure 1c, Supplementary Figure 10).

### Imaging neural activities over the whole larval zebrafish brain at high spatiotemporal resolution

To exemplify the imaging performance of Confocal LFM in real biological samples, we performed functional imaging of calcium activities over the entire brain of a movement-constrained larval zebrafish. Limited by the maximum frame rate of 30Hz for the employed camera (5120 × 3840 pixels), we could image ⌀ 800 μm × 200 μm volume (85 million voxels) in zebrafish brain at 6 Hz.

Elimination of background offers two key advantages: increased sensitivity for calcium activity detection and reduced reconstruction artifacts. In conventional LFM, the detectable calcium activity of a cell is limited by the Poisson noise from the fluorescence of its surrounding cells because LFM captures projection views that integrate all fluorescence along different directions. Although various computational methods can extract in-focus signal from the background, high background still leads to high Poisson noise^35^. Given that a larval zebrafish head is 300μm thick, the reduced axial coverage of 45μm in this Confocal LFM implementation offered 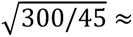 2.6 times increase in Signal to Noise Ratio (SNR). Even more importantly, the volume reconstruction algorithm will converge to a more accurate solution if limited number of projection views are captured from an effectively smaller volume. These effects can be clearly shown in Figure 2a, where spontaneous calcium activities were sequentially captured by LFM with and without confocal detection and with 67 ms time interval. Confocal LFM yielded sharper images with finer spatial details than conventional LFM. Detectable reconstruction artifacts in conventional LFM, such as several dark regions near the optical tectum, hindbrain and central line, were absent in Confocal LFM (Figure 2a, Supplementary Video 2). Bright neurons could be detected using both methods, but faint ones and their calcium activities were only detectable in Confocal LFM. Densely packed neurons (neuron 9-17 in Figure 2a) were much easier to resolve in Confocal LFM than in conventional LFM. This demonstrated significantly improved imaging resolution and sensitivity for calcium activity detection in living zebrafish brain.

**Figure 2.**
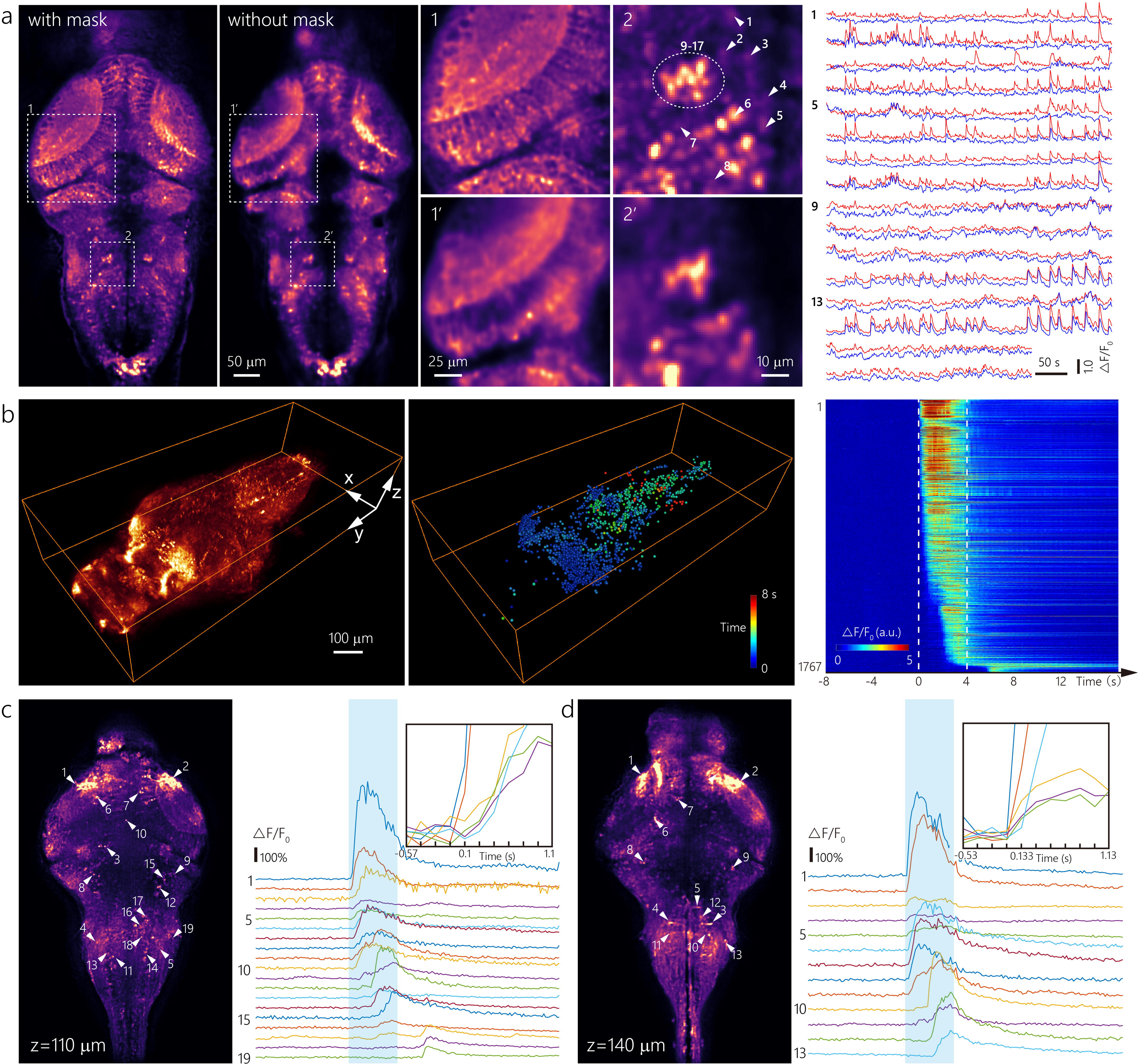
Whole brain functional imaging of neural activity in movement constrained larval zebrafish. (a) Maximum Intensity Projections (MIPs) over time of representing planes in reconstructed volumes obtained by Confocal and non-confocal LFMs when imaging spontaneous neural activity in larval zebrafish brain (left); Zoom-in views of example regions (middle); Neural activities of representing neurons obtained by Confocal (red) and non-confocal (blue) LFMs (right), (b) The MIP over time of calcium signals captured by Confocal LFM over the whole larval zebrafish brain when light stimuli was applied (left); Neural structures that were activated upon light stimuli are rendered as small spheres at their corresponding locations and colored by their onset time (middle). Neural activities of these neural structures ordered by their onset time (right). Dashed lines at 0 s to 4 s represent the start and end of light stimuli, (c) Representing neural structures at z = 110 *μ*m (left) and their fluorescent intensity traces (right). They were labeled in the order of their response onset times. Zoom-in view (upper right) of the fluorescent signals of first 6 neural structures, (d) Representing neural structures at z = 140 *μ*m (left) and their fluorescent intensity traces (right). They were labeled in the order of their response onset times. Zoom-in view (upper right) of the fluorescent signals of first 6 neural structures.

The high volumetric imaging speed can be useful for identifying the sequence of neural activation over the entire circuit. To illustrate this, we stimulated restrained larval zebrafish with a flash of light and observed a large population of neurons, starting from the optical tectum that mainly consists of visual sensory related circuits and ending at the hindbrain that was highly responsible for motor output, were strongly activated sequentially, which represented a classical sensory to motor transformation (Supplementary Video 3, 4). The calcium signal changes of the activated neurons could be extracted by combing both spatial and temporal features in the time series of imaging volumes, but the densely packed and structurally complicated dendrites that were also strongly activated could confuse the identification of somas of neurons in some brain regions. We extracted 1767 spatially non-overlapping footprints of neural structures that were activated upon light stimulus, as shown in Figure 2b, and quantified the sequence of their activation onset time. The densely packed neurites in the optical tectum were activated first and followed by somas at the outer edge of the optical tectum. Meanwhile, visible neural fibers starting from optical tectum were activated and signal was transmitted all the way down to lateral sides of the hindbrain. Several neurons located along this activated neural fiber tract and between optical tectum and hindbrain were then activated. They likely drove the neural activity near the middle line of the hindbrain later. This unique combination of the large imaging volume, high spatial and temporal resolution could provide new insight into the fast dynamics of the entire neural network in larval zebrafish brain.

### Whole brain functional imaging of neural activities in freely swimming larval zebrafish

The high volumetric imaging speed and epi-illumination configuration together make LFM a well-suited technique to image freely moving animals, which are of great interests in neuroscience because of the extended behavioral paradigms to understand various complex brain functions^19, 36–39^. In particular, neural activity in groups of neurons during larval zebrafish’s prey capture behavior, which is under extensive investigations^40–43^, has been recorded at relatively low spatial or temporal resolutions^31, 44, 45^.

To demonstrate the improved imaging performance of Confocal LFM in freely moving animals, we also combined it with a high speed 3D tracking system developed previously to image neural activity over the whole brain during larval zebrafish’s prey capture behavior (Supplementary Video 5). The fast tracking system employed a high speed camera and fast moving X-Y-Z stages to keep the larval zebrafish brain within the field of view. At the same time, behaviors of the zebrafish and the paramecia can be captured by another dedicated camera, as shown in Figure 3a-c. To avoid motion induced image blur, we finished scanning and camera exposure in 2 ms in each frame readout time of 33 ms (maximum camera frame rate of 30 Hz). The achieved equivalent instantaneous volume rate is 500 Hz.

**Figure 3.**
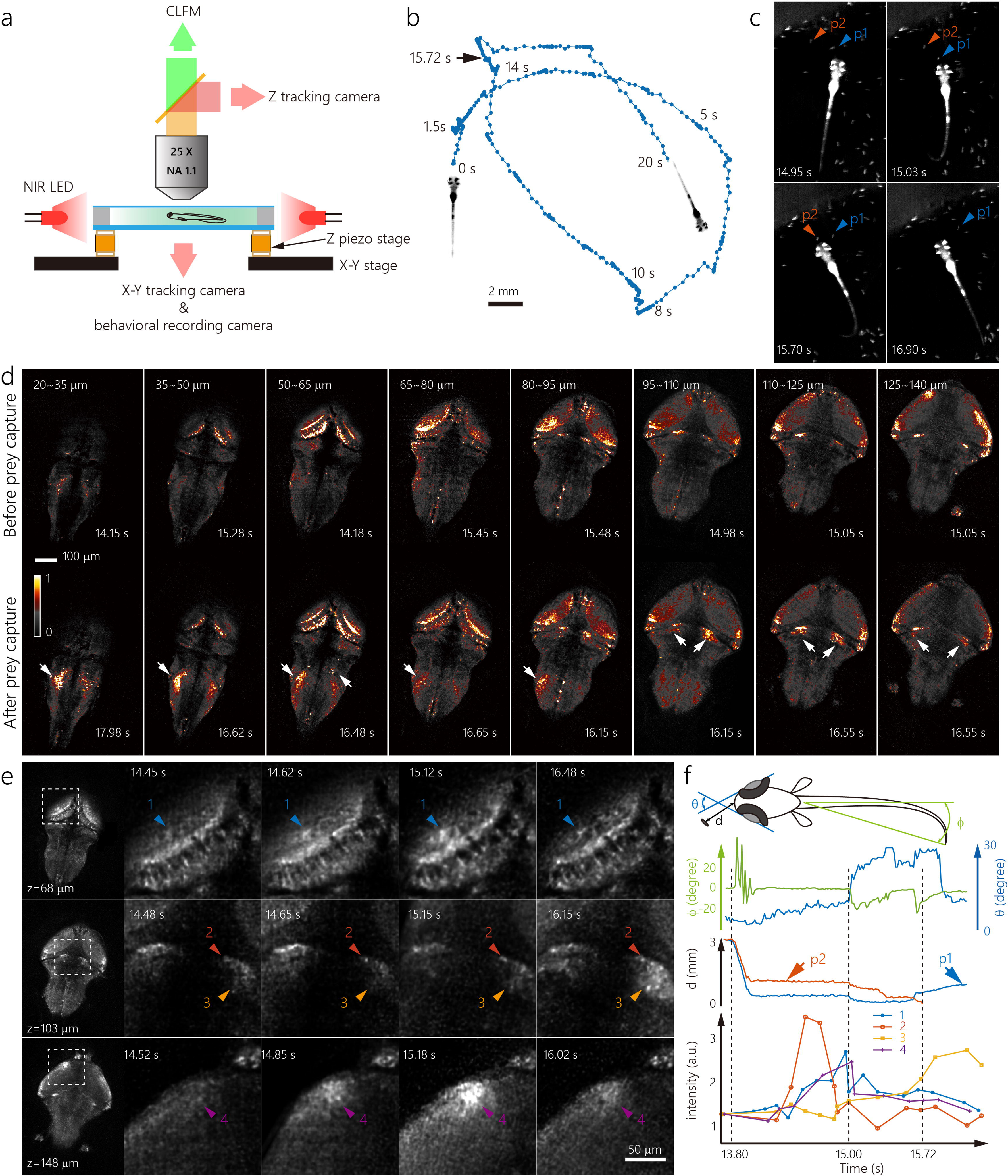
Tracking and imaging whole brain neural activity during larval zebrafish’s prey capture behavior. (a) Simplified schematics of the tracking and imaging system. NIR LED, Near InfraRed Light Emitting Diode, (b) The swimming trajectory of the larval zebrafish during the tracking and imaging. Successful capture of paramecium occured at 15.72 s. (c) Representing frames of behavioral recording during larval zebrafish’s prey capture. Two relevant paramecia were indicated by arrows, (d) Comparison of MIP images over indicated axial ranges at representing time points before and after successful capture of paramecium. Activated neurons and regions are indicated by arrows, (e) Zoom-in views of neural activities in several brain regions during larval zebrafish’s prey capture. Arrow 2 indicates a single activated neuron, (f) Kinematics of behavioral features and neural dynamics inferred from GCaMP6 fluorescence changes (Supplementary Figure 11) in four selected regions or neurons in (e) during the prey capture process. P1,2 refer to paramecium 1 and 2 in (c).

In an example prey capture event, the larval zebrafish initiated a prey capture represented by its eye convergence and J-turn pattern of its tail^40^ at around 15.03 s, but it failed to capture the first paramecium indicated by the blue arrow in Figure 3c. Right after the first try, it turned to another nearby paramecium indicated by red arrow in Figure 3c and ended with success. Activation of several brain regions upon its successful prey capture could be clearly identified as shown in Figure 3d. This result agreed well with previous observations^31^, but provided much higher resolution and sensitivity. For example, activation of several isolated neurons in the hindbrain could be clearly identified; two distinct groups of neurons near the cerebellum could be well resolved and only the group located posteriorly was strongly activated upon successful prey capture. More interestingly, we observed strong activation of several groups of neurons before the behavioral initiation of the prey capture. At ∼14.60 s, we observed neuralpills in dorsal optical tectum (indicated by arrow 1 in Figure 3e), a single neuron near the cerebellum (indicated by arrow 2 in Figure 3e), and a small region in ventral anterior optical tectum (indicated by arrow 4 in Figure 3e) were activated almost simultaneously. The larval zebrafish converged its eyes and made a J-turn 0.4 s later. Then calcium signal from these three regions started to decrease, indicating their possible roles in visual detection of the paramecium and initiation of the prey capture (Figure 3f). Only after the second successful prey capture, the calcium signal of a small group of neurons near the cerebellum (indicated by arrow 3 in Figure 3e) started to rise, which implied its distinct function from previous three groups. This example demonstrated that Confocal LFM could be successfully applied to image the whole larval zebrafish brain during its natural behavioral states and provided greatly improved spatial resolution and signal detection sensitivity.

### Volumetric imaging of neural activity with reduced background in scattering mouse brain

To further demonstrate the application of Confocal LFM to imaging larger brain, we recorded populations of neurons’ activities in awake mouse brain. Imaging into scattering tissue with conventional LFM is challenging because the ballistic light that survives from scattering and contributes to image formation decreases exponentially^13–15^. Meanwhile, the background outside the focusing volume becomes overwhelming as imaging depth increases. To better extract neural activities embedded in high background in LFM, different computational strategies making use of combined spatial and temporal features of activated neurons have been developed^27, 28^. Here, we show that Confocal LFM could improve Signal to Background Ratio (SBR) in raw data by physically rejecting background and increase the sensitivity to detect neural activities. This physical strategy is both compatible with and beneficial to various computational approaches and provides a greatly improved LFM platform for high speed volumetric functional imaging in thick tissues.

To achieve optimal imaging performance in mouse brain, we built another Confocal LFM with a spatial resolution of 4 × 4 × 6.5 μm^3^ over an imaging volume of ⌀ 800 μm × 150 μm (Supplementary Figure 12-15). We imaged calcium signals in the primary visual cortex of an awake mouse and did frame-by-frame unbiased volume reconstruction for flexible data processing and neural activity extraction. Three imaging volumes focusing at increasing depths were acquired to cover an axial range from the cortex surface to 400 μm deep. As shown in Figure 4a, both conventional LFM and Confocal LFM could image densely labeled neurons well near the surface of the mouse cortex, but Confocal LFM did better at suppressing background (Supplementary Figure 16) and therefore unveiled more weakly activated neurons than conventional LFM as imaging depth increased (Figure 4a, Supplementary Video 6). To quantify the improvement on SBR of Confocal LFM, we estimated the background using the low spatial frequency components in reconstructed raw images^46^ and the signal using the peak calcium activities of the 20 brightest neurons at each depth. As shown in Figure 4b, SBR increased when imaging depth went from ∼50 μm to ∼110 μm as more activated neurons appeared. After that, SBR started to decrease because scattering in tissue attenuated fluorescent signal exponentially as imaging depth increased. At depth greater than 300 μm, SBR estimation in conventional LFM was not meaningful anymore because the 20 brightest features detected were random fluctuations in the background rather than actual neurons. Confocal LFM out-performed conventional LFM over the entire depth range and provided about three times increase in SBR at ∼300 μm. More interestingly, Confocal LFM provided different SBRs at the same depth when the axial coverages were different. For example, Confocal 2 and Confocal 3 in Figure 4b represent two imaging volumes with different axial coverages of 150 μm — 300 μm and 250 μm — 400 μm, respectively. Confocal 3 provided better SBR than Confocal 2 at a presenting depth of 290 μm because the background noise from neurites above 290 μm are much stronger than that below it. This example showed that the background noise was the major limiting factor in functional imaging in thick tissue and the proposed confocal detection scheme could effectively suppress it in LFM imaging.

**Figure 4.**
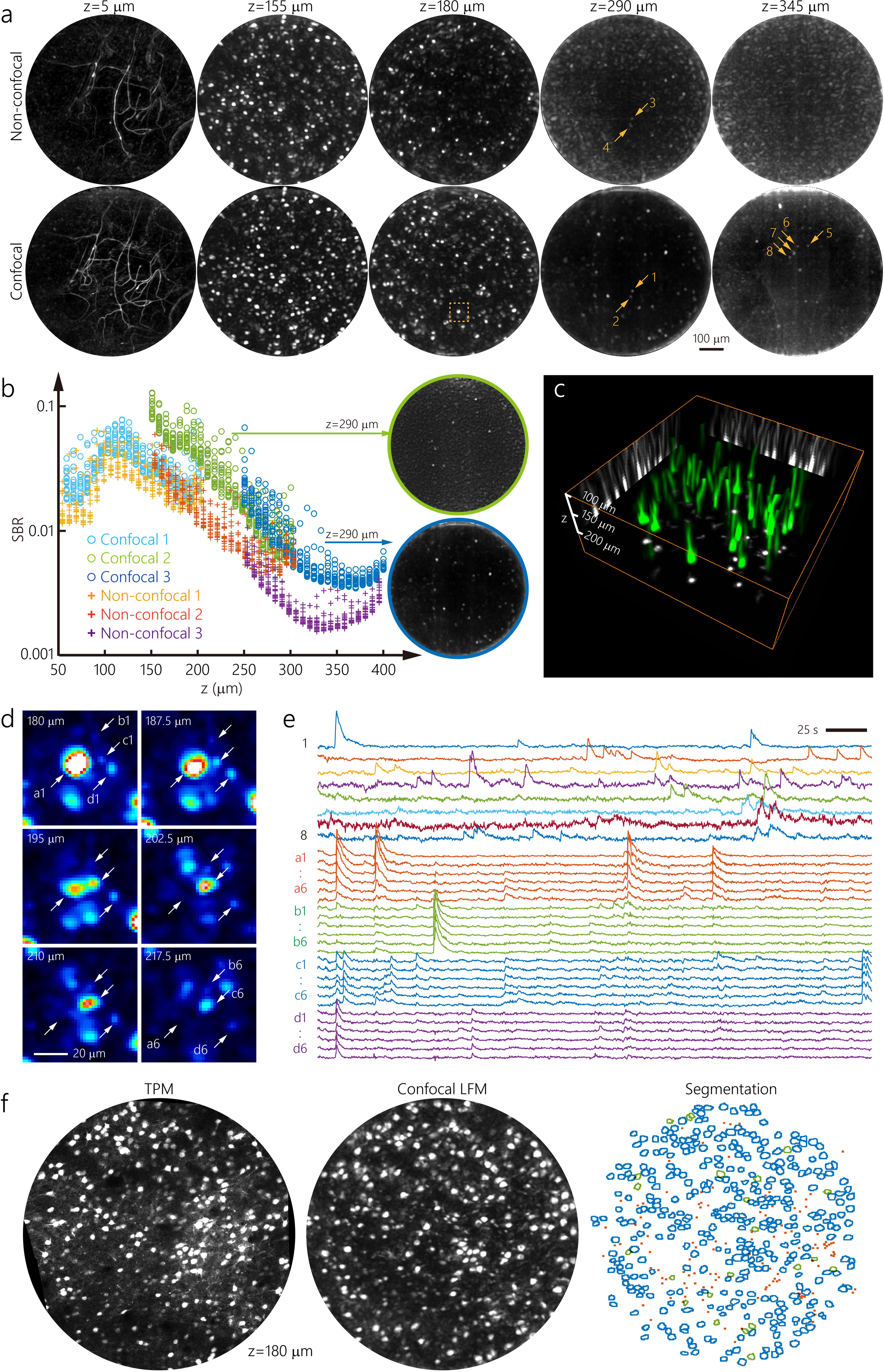
Volumetric functional imaging of neural activity in awake mouse brain. (a) Comparison of imaging results obtained by non-confocal and Confocal LFMs at representing depths in mouse cortex labelled with GCaMP6. Displayed images are standard deviations over time of background subtracted fluorescent images (Methods), (b) Experimentally measured SBR in Confocal and non-confocal LFMs at different depths. Confocal 1∼3 and Non-confocal 1∼3 indicate six different imaging trials carried out with Confocal and non-confocal LFMs and at different depth. Each data point represents SBR measured on one neuron. Top 20 brightest neurons are plotted at each depth in Confocal and Non-confocal cases. Insets with green and blue edges are imaging results processed in the same way as in (a) and obtained at z = 290 *μ* m in Confocal 2 and 3, respectively. (c)Top 50 longest neural structures identified by combing spatial footprints that were adjacent and functionally correlated, as found by CNMF-E at different z planes, (d) Zoom-in views of the indicated small region in (a) and its neighboring z planes. a1-a6, b1-b6, c1-c6, di-d6 are neural structures from four different neurons at increasing depths, (e) Neural activities of representing neural structures in (d) extracted by CNMF-E. (f) Comparison of MIP images over a volume of 20 *μ* m thick obtained by TPM (left) and Confocal LFM (middle). Time series of 500 volumes obtained by Confocal LFM were processed in the same way as that in (a) to generate the displayed image. Neural structures in both images were manually identified for comparison (right). Blue circles indicate neurons that could be found in both Confocal LFM and TPM images; Green circles indicate neurons could only be found in TPM image. Red dots were dendrites found both in Confocal LFM and TPM images.

The combined high resolution and sensitivity allowed us not only to extract calcium signal changes from activated neurons, but also to explore their structural features that could potentially be useful for confirming their identities. To do so, we extracted time series of different z planes from frame-by-frame reconstructed imaging volumes and identified spatial footprints of activated neural structures using CNMF-E^46^, a state-of-the-art constrained non-negative matrix factorization based method^47^, on each z plane independently. Because CNMF-E employed a ring model for background estimation and preferentially suppressed laterally extending neurites, the algorithm could detect cross-sections of vertical neural structures. By combing adjacent and functionally correlated neural footprints at different z planes, we could correctly detect trunks of pyramidal neurons’ epical dendrites, determine the correct locations of their somas and differentiate them from passing-by fibers (Figure 4c, Supplementary Video 7). This strategy worked even in some densely labeled regions, as shown in Figure 4d. Bright neuron somas and passing-by fibers as close as 8 μm could be well resolved. The cross-sections of continuous fibers can be correctly extracted as individual spatial footprints at different depth and their functional traces were highly correlated (Figure 4e, Supplementary Figure 17). To further confirm the imaging performance of Confocal LFM in mouse brain, we imaged the same populations of neurons by Confocal LFM and Two Photon Microsope (TPM). Although the brightness of the same neuron recorded in two different imaging modalities were different as they were imaged at different times, the spatial footprints of these neurons were highly correlated, as shown in Figure 4f. At a representing plane ∼180 μm below the cortex surface, Confocal LFM successfully detected 94% of neuron somas (362 out of 385) found in TPM. Meanwhile, we identified 119 cross-sections of neurites that could be resolved both by Confocal LFM and TPM in this plane. This high speed volumetric imaging and reliable extraction of neural structures and activities achieved in Confocal LFM only required very low laser dosage (about 1 mW over the entire imaging volume, Supplementary Table 2), therefore functional recordings over tens of thousands of volumes each day (Supplementary Figure 18) and over multiple days are feasible.

### Imaging and tracking circulating blood cells in awake mouse brain

Capturing fast dynamics in circulation system at microscales can help to provide insights into its developmental processes and mechanisms of disease states^48^. Various techniques have been developed to image beating heart in larval zebrafish^19, 30, 48^, but it is challenging to extend these techniques to image deep into large and scattering tissues, such as mouse brain. Here, we show that Confocal LFM could image fast circulating blood cells in thousands of blood vessel branches simultaneously and 600 μm deep in the awake mouse brain, which represented more than 100 times increase in throughput than conventional imaging methods^49^.

We labeled part of the circulating blood cells with a deep red fluorescent dye^50^ in awake mouse brain and imaged them at 70 Hz. Due to the reduced scattering at longer wavelength, we could image labeled blood cells up to 600 μm deep into the mouse cortex (Figure 5a, Supplementary Video 8). By tracking the fluorescent signal change along each blood vessel and over time, we could obtain circulating blood cells’ trajectories and trace them with high spatial and temporal precisions in 3D, as shown in Figure 5c. The speed of blood circulation, as manifested by the blood cells’ movement, was constantly changing due to either stochastic processes or biological meaningful processes^49^, such as blood vascular coupling (Figure 5d, Supplementary Video 9, 10). Moreover, blood cells did not keep constant traveling speeds across the vessels possibly due to their geometrical variations (Figure 5d). We analyzed 2554 vessel segments (∼80.4% of total blood vessels) in five imaging volumes ranging from 0 to 550 μm deep and tracked 61059 blood cells in five trials. These detailed measurements on each vessel were combined to provide an averaged estimation of the blood circulation dynamics over the entire imaging volume (Figure 5e). Furthermore, we could explore the hydrodynamic coupling between different blood vessel branches by doing clustering analysis based on the instantaneous velocities of the blood flow in them. For example, we divided the blood vessels at depth between 100 μm ∼ 250 μm into three different groups (Figure 5f). The average blood flow velocity in cluster 1 was more likely undergoing random changes. Cluster 3 contributed to a sudden rise in total averaged velocity at time 3 s. Cluster 2 and 3 together contributed to the velocity peak around 12 s. The spatial distributions of blood vessels in these three groups turned out to be intermingled.

**Figure 5.**
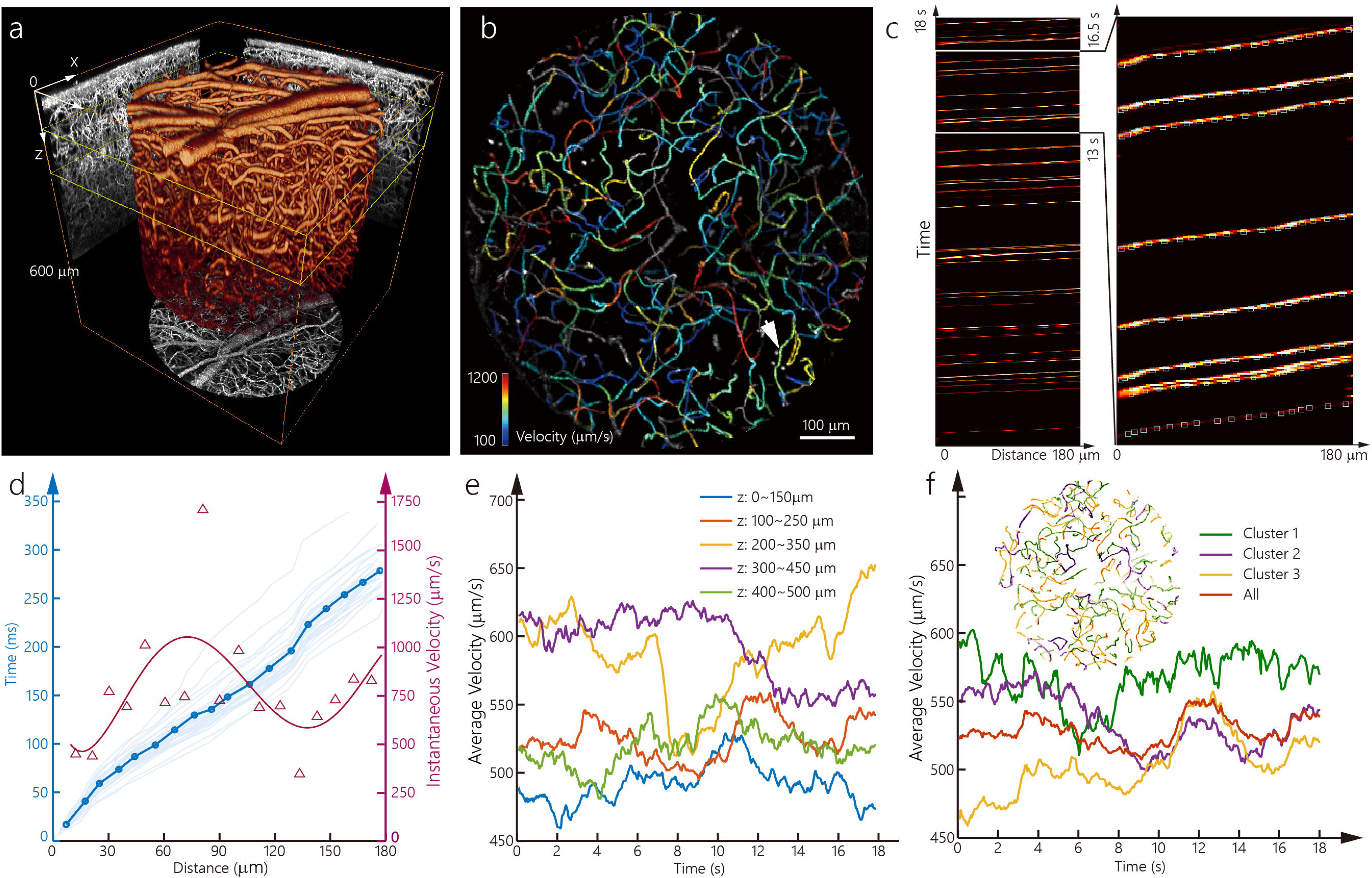
Imaging and tracking circulating blood cells in awake mouse brain. (a) volume rendering of vascular network obtained by averaging imaging volumes of circulating cells captured by Confocal LFM over time. Six time series of volumes imaged at six different depths are stitched together as one volume of more than 600 *μ*m in z (Supplementary Video 8). (b) MIP of the vascular network at depth between 100 *μ*m and 250 *μ*m. Blood vessel branches are color coded based on the average velocities of blood cells in them. Vessels colored in gray indicate failures in estimating their speed due to several reasons (Methods), (c) An example of cell trajectories captured on one representing blood vessel branch indicated by white arrow in (b). Each traveling blood cell can be identified as a continuous bright trajectory along the vessel. Their trajectories can be tracked precisely in time and space, as indicated by small squares on them in the zoom-in view (right), (d) Different cell trajectories are aligned in time and space showing the varying velocities between them (light blue). The average of these trajectories (dark blue) indicates non-constant traveling speed along the blood vessel. Instantaneous velocities along the vessel can be calculated by differentiating the averaged trajectory with respect to time (triangles in dark red). Solid line in dark red is a polynomial fit of it. (e) The average of instantaneous velocities over all the blood vessels in the imaging volumes change over time. Different curves refer to different experimental trials measured at different depths, (f) The average velocities over three different clusters of blood vessels in the depth range of 100*μ*m ∼ 250 *μ* m. Inset shows blood vessels color coded by their clusters.

## Discussion

Similar to confocal microscopy that greatly improves over wide-field microscopy the three dimensional fluorescence imaging capability in large tissues, Confocal LFM effectively reduces background in conventional LFM and provides greatly enhanced flexibility and imaging performance deep in the brain. These advantages come without compromising its characteristic high volumetric imaging speed. Although the scanning of vertical excitation light sheet in Confocal LFM to cover a volume is not instantaneous, the camera frame rate rather than the scanning speed, which can ramp up to kilohertz, limits the practical volume rate. In fact, we could achieve much higher instantaneous volume rate by making scanning speed faster than camera frame rate. This can be critical in many imaging applications. For example, to avoid motion induced image blur, we finished scanning and camera exposure in 2 ms in each frame readout time of 33 ms (30 Hz frame rate) when imaging freely behaving larval zebrafish. The achieved equivalent instantaneous volume rate is 500 Hz. Without high instantaneous volume rate, high quality imaging in such situation would not be possible. We also applied the same strategy to get blur-free images of circulating blood cells by setting instantaneous volume rate as high as 333 Hz (3 ms exposure time) when camera ran at 70 Hz.

Besides confocal detection, selective plane illumination is another effective method to suppress background. In particular, oblique plane illumination and imaging through a single objective preserved the epi-illumination configuration and greatly enhanced its compatibility with biological studies^19, 51^. Introducing this strategy to conventional LFM could also be attractive, but the large incident angle of the oblique illumination beam could make it difficult to maintain its beam shape in scattering tissue and reduce the efficiency of background suppression.

The practical performance of LFM heavily relies on characteristics of employed image sensors. More specifically, LFM requires high speed, low noise, large number of pixels, and square-shaped image sensor. However, commercially available sensor chips of this type are usually of industrial grade and have higher readout noise than scientific grade cameras. We employed an industrial grade camera for its large pixel number in Confocal LFM optimized for zebrafish imaging, but we expect much better imaging performance, particularly on picking up weak signals, if a scientific grade camera with similar number of pixels can be used.

We demonstrated the potentials of Confocal LFM in applications such as high spatiotemporal resolution imaging of whole brain neural activity in freely behaving larval zebrafish. Because of the improved resolution and sensitivity, we could identify behaviorally correlated individual neurons’ activities during larval zebrafish’s prey capture. However, registration of every single neuron and reliable extraction of their activities over long period of time on freely swimming larval zebrafish are still challenging because the animal’s body, including the brain, could undergo large deformation and compromises the registration even with the improved spatial resolution. Reducing labeling density slightly or specifically labeling certain groups of neurons can be practical strategies if highly reliable extraction of single neurons’ activities over large population and long period of time is critical in imaging freely moving animals.

Applying Confocal LFM to calcium imaging in densely labelled mouse brain achieved better SBR and larger penetration depth at the cost of reduced imaging volume coverage. This trade-off can potentially be circumvented by introducing multi-focal microscopy into Confocal LFM, as demonstrated in XLFM^31^. In post data processing, we adopted frame-by-frame reconstruction strategy for unbiased structural estimation and applied CNMF-E separately on different z planes. This unbiased volume reconstruction helped to clearly differentiate somas from passing-by fibers and showed highly correlated imaging results with that obtained by TPM. If unbiased structural information is not critical, several computational methods developed for conventional LFMs^27, 28^ could be readily applied to Confocal LFM to extract neural activities from somas and lead to better performance than conventional LFM. Ideally, a new algorithm that not only combines spatial and temporal features to extract neural activity, but also provides unbiased 3D structural reconstruction at the same time in LFM would be highly valuable. Additionally, the demonstrated capability to correlate densely labelled neurons captured in Confocal LFM to that imaged in TPM could open ways to combine the strengths of speed and resolution in these two imaging techniques. For example, we could identify neurons that are interesting for certain experimental purposes in large populations of neurons imaged by Confocal LFM first and then characterize dendrites of these neurons at high resolution using TPM to get more detailed structural or functional information.

Finally, we showed that circulating blood cells could be well tracked using Confocal LFM over a large volume and at 600 μm deep into the mouse brain. It provides a unique way to precisely quantify the blood flow dynamics and coupling between different vessels in a complicated 3D vascular network. Furthermore, this system can be readily combined with volumetric functional imaging to explore the coupling between neural activity and vascular system, which underlies a range of noninvasive neuroimaging techniques and are of great interests and importance to neuroscience^49, 52^.

### Data availability

The data that support the findings of this study are available from the corresponding authors upon request.

### Code availability

Custom-written software code for system coordination and data acquisition (Labview) and volume reconstruction in Confoal LFM (Matlab) is available upon request.

## Acknowledgements

K.W. acknowledges supports from National Key R&D Program of China (2017YFA0700500), Strategic Priority Research Program of the Chinese Academy of Sciences (XDB32030200), International Partnership Program of Chinese Academy of Sciences (153D31KYSB20170059), Shanghai Municipal Science and Technology Major Project (2018SHZDZX05), NSFC (31871086) and China Thousand Talents Program. We thank Dr. Jie He for his support on zebrafish handling and helpful discussions.

## Author Contributions

Z.Z. and L.C. built Confocal LFM. Z.Z. wrote the code for image reconstruction. Z.Z. and T.Z. performed zebrafish related experiments and data analysis. L.B. and P.Y. performed experiments on mouse. L.B. performed data analysis on calcium imaging on mouse. L.B. and W.S. performed data analysis on blood flow in mouse brain. Z.Z., F.L. and J.D. designed zebrafish behavioral experiments. K.W. conceived and led the project. Z.Z., L.C., L.B. and K.W. wrote the manuscript with input from all authors.

## Confocal LFM for zebrafish brain imaging

The imaging system (Figure 1a, Supplementary Video 1) was modified from our previously reported XLFM^31^. In the laser excitation light path, a 473 nm laser (cnilaser MBL-N-473-1.5W) was reflected from a scanning galvo (SINO-GALVO JD1105, not shown in Figure 1a) to direct laser beam into and out of microscope rapidly. It was then expanded to 10 mm diameter collimated beam and focused into a line by a cylindrical lens (f= 1000 mm, Thorlabs LJ1516RM, not shown in Figure 1a). This line was then conjugated to the mask by an achromatic lens (f= 150 mm, Thorlabs AC254-150-A, not shown in Figure 1a) and a telecentric lens L3 (f= 35.1 mm, Daheng Optics GCO-2303). A scanning galvo (SINO-GALVO JD2206) was placed between these two lenses to scan the focused laser line (in × direction) along y direction. 15 masks were mounted to a direct-drive rotation stage (YIDING DF1A-004G) to scan at a constant speed (120 rpm at the camera frame rate of 30 Hz). Glass plates of different thicknesses were directly attached to their corresponding masks using UV-curing adhesive (Thorlabs NOA61). During imaging, the galvo scanned the focused laser line at the same speed as the rotating mask so that laser beam could go through thin slit in the center of each mask (Supplementary Figure 1b, Supplementary Video 1). Additionally, the galvo was conjugated to the central micro-lens in the array such that the laser beam going through the central slit on the mask will always be directed to the central micro-lens in the array during scanning. The micro-lens array (Supplementary Figure 1c) was conjugated to the back pupil of the imaging objective (Nikon CFI75 Apo 25XC W, NA 1.1) by two identical achromatic lenses L1 and L2 (150 mm, Thorlabs AC508-150-A). The fluorescence excited by the vertical blue light sheet is collected by the imaging objective and relayed to micro-lens array. The micro-lens array consisted of 25 micro-lenses (f= 38 mm, 3 mm in diameter) that formed 25 fluorescence images on the scanning mask. These images were spatially filtered by 25 apertures, including the central slit, on the masks and then conjugated to the camera (Ximea CB200MG-CM, 5,120 × 3,840 pixel) by L3 and L4 (f= 54.37 mm, Daheng Optics GCO-2302). A dichromic mirror D1 (Semrock FF496-SDi01-25×36×2.0) and a band pass filter F1 (Semrock FF03-525/50-32) were employed to spectrally separate the blue laser and the fluorescence. To synchronize the galvo and rotating masks, we designed an extra aperture, or trigger slit, on each mask (Supplementary Figure 1b) such that a 450nm laser beam (Thorlabs CPS450, Supplementary Figure 2) focused at the slit could pass the mask and hit a light detector behind (Thorlabs PDA36A2, Supplementary Figure 2) when the rotation stage was at the desired location. During scanning, the stage with 15 mounted masks rotated at a constant speed. Every time the light detector detected a pulse from 450 nm laser, it sent triggers to turn on the 473 nm laser and start the galvo scan and camera exposure (Supplementary Figure 2). Because the spacing between the trigger slit and the central slit could be established very precisely when manufacturing the mask, the synchronization of the system was very accurate.

## Confocal LFM for mice brain imaging

Different from imaging larval zebrafish, imaging blood flows in mouse brain required very high speed and left no room for axial scanning. Therefore, we built a system without fast axial scanning capability, i.e., the glass plates of various thickness, as shown in Supplementary Figure 12. The system employed two pairs of achromatic lenses (L1, f= 150 mm, Edmund Optics #49-391; L2, f= 160 mm, Edmund Optics #49-382; L3 & L4, f= 100 mm, Thorlabs AC254-100-A) to conjugate the back pupil of the imaging objective (Olympus XLPLN25XWMP2, NA 1.05) to a scanning galvo (G1, SINO-GALVO JD2808) and a customized micro-lens array, which consisted of 37 micro-lenses (f= 26 mm, 1.3 mm diameter). The excitation laser (Coherent OBIS 488-100-LS or cnilaser MLL-III-635L-100mW) were shaped into rectangular 1.3 × 20 mm^2^ beams before introducing into the system by a dichroic mirror (D1, Semrock Di01-R488/561 or Optolong BP-DM11A). L4 focused the laser beam into a line and G1 scanned this line to cover a volume under the objective during scanning. The excited fluorescence was de-scanned by G1 and formed 37 images after micro-lens array. These images were filtered by an optical mask placed at the focal plane of the micro-lens array. An additional pair of telecentric lenses (L5 & L6, f= 35.1 mm, Daheng Optics GCO-2303) conjugated the mask to the imaging camera (Hamamastu ORCA-Flash4.0). Because in this configuration G1 also de-scanned the fluorescent images, a second scanning galvo mirror (G2, SINO-GALVO JD2808) was placed between two telecentric lenses and conjugated to objective’s pupil plane to re-scan the images. The two galvo mirrors were synchronized to ensure correct de-scan and re-scan. The range of scanning angles of them were carefully determined experimentally such that the scanned distance in y in the sample plane matches that in the scanned image on the camera. Band-pass filters (Semrock FF03-525/50 or Optolong EM692/40) were inserted right before the camera to block excitation lasers.

## Design and assembly of micro-lens array, masks and glass plates

Micro-lens arrays were manually assembled by mounting customized micro-lenses onto customized lens housings (Supplementary Figure 1c). The micro-lenses in an array were designed to have identical NAs of 0.13 and 0.07 in Confocal LFMs optimized for zebrafish and mice brain imaging, respectively. To better match masks for these micro-lens arrays, we characterized their properties experimentally in the following way. The detection path up until the mask was aligned and a camera was placed at the focal plane of this micro-lens array, i.e., in the place of the mask. A fluorescent bead was placed under the objective and near the center of the field of view, and excited by a laser beam from the bottom. Camera recorded two images of the bead when it was placed at the top and bottom of the designed focusing range. Connecting these two images of the bead recorded at different axial positions by lines to generate a binary image. Convolving this image with another line-shaped binary image representing the shape of the focused excitation beam resulted in the final mask design (Supplementary Figure 3). The right thicknesses of glass plates providing required axial shifts were determined experimentally by gradually accumulating 1 mm thick glass plates between the camera and the micro-lens array until the desired thicknesses were reached. Glass plates with right thicknesses and anti-reflection coatings were custom made based on these measurements.

## PSF measurement and image reconstruction in Confocal LFM

The measurement of PSF and reconstruction of 3D volumes were similar to that described in our previous work^31^. In short, a fluorescent bead of 1 μm diameter (ThermoFisher F8888) in the center of the field of view was scanned axially under the imaging objective and 3D imaging stacks were collected by Confocal LFMs at the same time. 15 PSFs measurements were acquired, one for each of the 15 masks in Confocal LFM optimized for zebrafish imaging. These 15 PSFs were measured on the same fluorescent beads in a single axial scan. Therefore, the small image shifts caused by different sets of masks and glass plates would be automatically compensated by incorporating their corresponding PSFs in volume reconstructions. Due to the same reason, the continuity along z axis and between neighboring volumes was guaranteed automatically. The voxel sizes of the PSFs measured in the two Confocal LFMs were 1 × 1 × 1.5 μm^3^ and 1.9 × 1.9 × 2.5 μm^3^, respectively.

In Confocal LFM optimized for zebrafish imaging, optical aberrations of the system was evident (Supplementary Figure 7). Therefore, we carried out aberration corrections in the volume reconstruction, as described in the following section. The reconstruction algorithm was based on Richardson-Lucy deconvolution and iteratively refined 3D estimation of the object. It usually took 30-50 iterations to converge to stationary results. We employed a CUDA program to do imaging reconstruction on a desktop computer (CPU: Intel i9-9900K, RAM: 32 GB, GPU: NVIDIA GeForce RTX 2080 Ti). It took about 80 s and 40 s to reconstruct one volume in Confocal LFMs optimized for zebrafish and mice brain imaging, respectively. Because the frame-by-frame reconstruction could be carried out in parallel, the reconstruction speed could be easily increased by scaling up the number of employed GPUs.

## Correction of system aberration

The system aberration did not distort the PSF, but it caused different field distortions in the images formed by different micro-lenses and resulted in low quality reconstructions (Supplementary Figure 7). To calibrate the system aberration, we imaged a layer of sparse fluorescent beads near the focal plane. The position of bead *i* in the image formed by micro-lens *j* on the camera could be localized as (*x_ij_*, *y_ij_*). Assuming the center of the field of view of the micro-lens *j* was (*x*_0*j*_, *y*_0*j*_), then the relative coordinates of the beads in sub-images formed by each micro-lens could be calculated as (*x_ij_* – *x*_0*j*_, *y_ij_* – *y*_0*j*_) = (Δ*x_ij_*, Δ*y_ij_*). Because of the field distortion, (Δ*x_ij_*, Δ*y_ij_*) ≠ (Δ*x_ik_*, Δ*y_ik_*) when *j* ≠ *k*. We chose the central micro-lens (*j* = 0) as reference and the goal is to find the transform for each micro-lens such that *f_j_* (Δ*x_ij_*, Δ*y_ij_*) = (Δ*x_i_*_0_, Δ*y_i_*_0_). In practice, we found that quadratic polynomial surface could fit this transform well (Supplementary Figure 8). Then we could match the images formed by different micro-lenses by applying the calibrated coordinates transforms and image interpolations. We incorporated these transforms into the reconstruction algorithm and experimentally confirmed that overall reconstruction quality was improved significantly (Supplementary Figure 7-9).

## Resolution characterization in Confocal LFM

The overall resolution of the system was determined by two factors: (1) the diffraction limit of each micro-lens and (2) the reconstruction algorithm that combined information from all micro-lens.

1. The diffraction limit of each micro-lens could be directly measured from PSFs produced by them (Figure 1b). Theoretically, the resolution limit of the entire system is determined by the micro-lens with best resolution. Therefore, we could experimentally determine that the diffraction-limited resolution of the system was about 2.13 × 2.13 × 2.5 μm^3^, as found on a representative outermost micro-lens (Supplementary Figure 6).
2. The overall resolution was also determined by reconstruction because the above diffraction limited resolution could be achieved only if the reconstruction could properly combine information from different micro-lenses together without contradictions among them. This was achieved after correcting system’s aberration (Supplementary Figure 7). We further characterize the performance of reconstruction by imaging fluorescent beads sparsely spread on a glass slide. The measured FWHM size of individual isolated beads after reconstruction was on average 1.2 × 1.2 × 1.7 μm^3^. Although this result was smaller than the diffraction-limited resolution, it did not suggest super-resolution as Richardson-Lucy based deconvolution methods tend to restore an isolated point-like object to a smallest possible spot. Therefore, this result only indicated that the best achievable resolution in final reconstruction under the condition of current noise level and aberration correction precision was about 1.2 × 1.2 × 1.7 μm^3^. Because this size was smaller than diffraction limit, the reconstruction was optimal and preserved diffraction-limited performance.

Based on experimental characterizations of above two factors, we concluded that the Confocal LFM optimized for zebrafish imaging provided diffraction-limited resolution of about 2.13 × 2.13 × 2.5 μm^3^ over the entire imaging volume of ⌀ 800 μm × 200 μm. Based on similar measurements and arguments, the Confocal LFM optimized for mice brain imaging provided diffraction-limited resolution of ∼ 4 × 4 × 6.4 μm^3^ over the entire imaging volume of ⌀ 800 μm × 150 μm.

## Comparison of Confocal LFM with non-confocal LFM

To compare the performance of Confocal LFM and non-confocal LFM when imaging zebrafish brain, we replaced one mask with a plain glass plate of similar thickness on the rotating stage and collected images through this glass plate and the mask next to it. Because the camera was run at 15 Hz, the time delay between these two images was 67 ms. Two different PSFs for these two cases were measured as described as above and image reconstructions were carried out using their corresponding PSFs. Similarly, the comparison of Confocal LFM and non-confocal LFM in mouse brain imaging was achieved by keeping and removing the mask behind micro-lens array.

## Imaging fluorescent beads in agarose

Fluorescent beads of 0.5μm in diameter (ThermoFisher F8888) were mixed with 1.2% low-melting agarose (Sigma-Aldrich A6877) before it turned into gel. A thin layer of this mixture was poured on to a glass slide and covered by a cover glass. The resulting thickness of the agarose between glass slides and cover glass was about 1 mm.

## Zebrafish preparation

All experimental procedures were approved by the Animal Care and Use Committee of the Institute of Neuroscience, Chinese Academy of Sciences. Adult zebrafish (*Danio rerio*) were maintained at the National Zebrafish Resources of China (NZRC, Shanghai, China) with an automatic fish housing system (ESEN, China) at 28°C on a 14/10 light/dark cycle following a standard protocol^53^. Embryos were raised in 10% Hank’s solution, which consisted of (in mM): 140 NaCl, 5.4 KCl, 0.25 Na_2_HPO_4_, 0.44 KH_2_PO_4_, 1.3 CaCl_2_, 1.0 MgSO_4_ and 4.2 NaHCO_3_ (pH = 7.2).

Transgenic zebrafish larvae (Tg(HuC:GCaMP6s)) were in *albino* background. All experiments were performed on larvae aged at 4-9 days post-fertilization (dpf) at 26-27 °C. For the imaging of a restrained larva, 4-6 dpf zebrafish were paralyzed with α-bungarotoxin (100 μg/ml, Tocris) for 5-15 min and embedded in 1.2% low-melting agarose (Sigma-Aldrich).

## Visual stimulation on movement restraint zebrafish

Light from a 415 nm LED was focused by a lens into a spot of 2.5 mm in diameter and positioned on the left anterior part of the movement restrained larval zebrafish brain. The power of the beam was about 1 mW.

## Zebrafish tracking system

The 3D tracking system was similar to our previous demonstration^31^. In brief, a fast tracking camera (2 ms exposure time, 300 fps or higher, Basler aca2000-340kmNIR, Germany) captured images of scattered NIR LED light from larval zebrafish. A customized LabVIEW program then extracted orientation direction and centroid of the zebrafish from these images at high speed (250 Hz). A Proportional–Integral–Derivative (PID) based control algorithm drove a two-axis direct drive stage (Newport ONE-XY60) to compensate the in-plane movement of larval zebrafish in arena. The tracking in z was achieved by a customized detector consisting of a micro-lens array conjugated to the back pupil of the imaging objective and a camera (100 fps or higher, Basler aca2000-340kmNIR, Germany) placed at its focal plane. The depth information was extracted by calculating the change of distances between images captured by different micro-lenses. The offset between the actual axial position of the fish head and the set point was fed into the PID to drive a pair of piezo actuators (Figure 3a, P06.X500R, Harbin Core Tomorrow Science & Technology Co., Ltd.) to compensate axial movement of larval zebrafish.

## Zebrafish prey capture behavior

In each experimental trial, one single 7-9 dpf zebrafish with 10% Hank’s solution was transferred into a customized chamber (42 mm in diameter, 0.85 mm in depth) together with 100-200 paramecia. The chamber was then covered by a coverslip (0.17 mm in thickness) for imaging. To increase the occurrence of prey capture event, zebrafish larva was food restrained for at least 12 hours before experiments^54^.

## Mice care and imaging

All experimental procedures were approved by the Animal Care and Use Committee of the Institute of Neuroscience, Chinese Academy of Sciences. All mice were housed in individual cages in 12hour light and 12hour dark room without water or food restraint during the whole experiment procedure.

For functional imaging of neural activity, 8-week-old male C57BL/6J or male SST-Cre (Jax 013044) mice were used. Mice were anesthetized by isoflurane during the surgery (3% for induction, 1%-1.5% during the surgery) and fixed onto the stereotaxic apparatus once they were fully sedated. Eyes were covered with erythromycin ointment to prevent drying and glare. The scalp was then removed and 3% hydrogen peroxide were applied on the skull to remove remnants. Primary visual cortex (V1) was labeled by stereotactic technique (AP: 3.8 mm ML: 2.6 mm from Bregma site). A customized titanium headpost was stuck to skull with glue (Loctite 495). A craniotomy (∼3 mm diameter) was administrated with skull drill. Craniotomy region was submerged in sterile saline during the drilling process to cool down the surface of brain. After removing the skull piece, virus mixture (AAV2/8-hSyn-Cre-WPRE-pA and AAV2/9-hSyn-FLEX-GCaMP6s-WPRE-pA, TaiTool BioScience) was injected in V1 with WPI injection system (75 nl/site, 3-5 sites/mouse). After injection, a cranial window consisting of three pieces of cover glass (one 5 mm in diameter and two 3 mm in diameter, 0.17 mm thick) sealed together by UV curing adhesive was implanted on the craniotomy region. Skull was dried with sterile sponges. Cranial window was attached to skull with glue (Loctite 495) and permanently secured with dental acrylic. Mice were allowed to recover for at least one week before imaging.

For circulating blood cell imaging, 6-8-week-old male C57BL/6J mice with cranial windows, as described above, were used. We adopted the Red Blood Cell (RBC) staining protocol as previously described^55^. In brief, a total of 150 μL blood was drawn from orbital vein (ipsilateral eye from imaging region) when mice were slightly anesthetized with 2% isoflurane. Blood was added into 1.4 mL blood plasma buffer (BPB, in mM: 128 NaCl, 15 glucose, 10 HEPES, 4.2 NaHCO_3_, 3 KCl, 2 MgCl_2_, and 1 KH_2_PO_4_, pH 7.4), gently mixed and centrifuged at room temperature at 1500 rpm for 5 minutes. Supernatant was discarded. RBC was re-suspended with dye mixture (1.4 mL BPB, 1.4 mL diluent C (Sigma CGLDIL), 1.2 μL DiD solution (DiD solid; Invitrogen D-7757, 0.1 mL ethanol dissolved)) and incubated in 37°C shaker for 5 minutes. 1.4 mL serum was then added and incubated for another minute. After incubation, the RBC suspension was centrifuged at 1500 rpm for 5 minutes and gently washed twice with 3.5 mL BPB and 350 μL serum. In the end, RBC was isolated by centrifugation at 1500 rpm for 5minutes and re-suspended in 200 μL BPB. The suspension was injected back into mice tail vein. About 10% of total blood cells were labeled in this process.

Before imaging, mice were slightly anesthetized with 2% isoflurane and head-fixed on the customized holder with body restricted in a plastic tube. The imaging window was cleaned with 75% ethanol. Mice were mounted on a 3-axis translation stage. After mice recovered from anesthetization, they were kept in the system for at least 15 minutes for habituation before imaging sessions.

## Registration of zebrafish brain

A reference brain was generated first by stitching multiple imaging volumes together based on the overlap regions between them. This was done semi-automatically. All imaging volumes were then registered to this reference brain in two steps. First, rotations along z axis were determined coarsely by maximizing the cross correlations between imaging volumes and the reference brain. Second, common structural features in reference brain and imaging volumes were extracted automatically and optimal affine transformations were determined based on fine registrations of these features. The registration results were manually inspected and failures in automatic registrations were replaced by manual registration results.

## Neural activity extraction in zebrafish and mouse brain

In movement-restrained larval zebrafish imaging experiments (Figure 2), time series of imaging volumes were registered first to cancel residual movement of the zebrafish. Then the correlation of time traces between each voxel and its neighboring 5 × 5 × 3 voxels, which roughly approximated the size of a neuron in larval zebrafish brain, were calculated to generate correlation images^56^. Local maximums in correlation images were then identified as centers of individual region of interest (ROI) for neural activity extraction. The ROIs were determined under two constraints: first, they had maximum size of 5 × 5 × 3 voxels; second, the minimum correlation values of pixels in the ROI had to be large than 70% of their maximums. To estimate the baseline fluorescence F_0_, we calculated the average of the entire time traces extracted from ROIs first. Time points when signal exceeded 20% of the calculated average were excluded and the average of the rest of the time points was used as the baseline F_0_.

In freely moving larval zebrafish imaging experiments (Figure 3), time series of imaging volumes were registered as described in methods above. The neural activity could not be directly extracted because the movement of zebrafish would cause intensity fluctuation in images even in silent neurons. The fluctuation was mainly ascribed to the body rotation during its swimming.

Fluorescence from neurons deep inside zebrafish had to go through different parts of the tissue to reach imaging objective when its body rotated. To quantify the neural activities in some regions that could be clearly identified by comparing with their relatively less activated surrounding regions, we calculated the intensity ratio between them (Supplementary Figure 11).

In head-fixed mouse imaging experiments (Figure 4), the frame-by-frame reconstructed imaging volumes were registered first using cross correlation. When registration became difficult for volumes captured at greater depth, we manually identified bright features as landmarks for registration. To increase SNR, voxels along z axis were binned by a factor of 3 and the original voxel size of 1.9 × 1.9 × 2.5 μm^3^ was increased to 1.9 × 1.9 × 7.5 μm^3^. In subsequent processing, individual z planes were extracted from the volumes and processed independently. We first estimated time-varying background by applying low-pass Gaussian filter (δ= 20 pixels) to each frame, which was similar to CNMF-E. Then the background was subtracted from each frame and positive constraint was applied to the result. Finally, standard deviation was calculated on the time series to generate images shown in Figure 4a. To extract neural activities, we feed raw data that was registered but not background subtracted to CNMF-E program we downloaded from https://github.com/zhoupc/CNMF_E. To merge the neural structures of the same neuron on different z planes, as shown in Figure 4e, we searched for spatial footprints identified by CNMF-E that were spatially close (the lateral shifts of spatial footprints’ peaks between neighboring z planes had to be smaller than 5.7 μm) and temporally correlated (Pearson correlation coefficient > 0.85).

## Analysis of blood flow in mouse brain

The time series of reconstructed imaging volumes were averaged over time first to get the 3D morphology of blood vessels. If sample movement was evident as blurry averaging results indicated, we registered the involved frames based on some blood cells that happened to be stuck in blood vessels. After getting a sharp reconstruction of the blood vessel network, we applied locally adaptive threshold to binarize the image. The blood vessels in the binary images were then further thinned down into lines to get skeleton of the network (Supplementary Video 10). To simplify the data processing, we excluded all branching points in the skeleton and divided the connected network into a list of unbranched vessel segments. For each vessel segment, we extracted time traces of fluorescent signal on each voxel and combined all of them sequentially in this segment to form an image (Figure 5c). The trajectories of blood cells can be identified as continuously ramping curves (Figure 5c). The precise locations of the cells at each time point were determined by centroid estimations. To quantify the changing blood flow velocity over time, we calculated the velocities of labeled cells averaged over the time in which they traveled across the vessel segment and linearly interpolated in between the time points when labelled cells were present in the vessels. In some blood vessels, the blood flows were too fast or the blood vessel segments were too short such that blood cells traveled through in less than 3 imaging frames and speed measurements could be inaccurate. In some other blood vessels, there were less than 5 blood cells traveling across during the entire imaging period. We excluded these vessel segments when calculating the average blood flow velocity over the volume. The clustering analysis of the blood vessels were based on the Pearson correlation of their time traces of velocities. The percentage of analyzed blood vessels were calculated as the ratio between total number of voxels in the analyzed vessel segments and the total number of voxels in the entire skeleton including branching points.

## References

1. Denk, W., Strickler, J.H. & Webb, W.W. Two-photon laser scanning fluorescence microscopy. Science 248, 73 (1990).

2. Katona, G. et al. Fast two-photon in vivo imaging with three-dimensional random-access scanning in large tissue volumes. Nat Methods 9, 201–208 (2012).

3. Kong, L. et al. Continuous volumetric imaging via an optical phase-locked ultrasound lens. Nat Methods 12, 759–762 (2015).

4. Nadella, K.M.N.S. et al. Random-access scanning microscopy for 3D imaging in awake behaving animals. Nat Methods 13, 1001 (2016).

5. Amir, W. et al. Simultaneous imaging of multiple focal planes using a two-photon scanning microscope. Opt. Lett. 32, 1731–1733 (2007).

6. Cheng, A., Gonçalves, J.T., Golshani, P., Arisaka, K. & Portera-Cailliau, C. Simultaneous two-photon calcium imaging at different depths with spatiotemporal multiplexing. Nat Methods 8, 139–142 (2011).

7. Prevedel, R. et al. Fast volumetric calcium imaging across multiple cortical layers using sculpted light. Nat Methods 13, 1021 (2016).

8. Yang, W. et al. Simultaneous Multi-plane Imaging of Neural Circuits. Neuron 89, 269–284 (2016).

9. Zhang, T. et al. Kilohertz two-photon brain imaging in awake mice. Nat Methods 16, 1119–1122 (2019).

10. Lu, R. et al. Video-rate volumetric functional imaging of the brain at synaptic resolution. Nat Neurosci 20, 620 (2017).

11. Song, A. et al. Volumetric two-photon imaging of neurons using stereoscopy (vTwINS). Nat Methods 14, 420 (2017).

12. Kazemipour, A. et al. Kilohertz frame-rate two-photon tomography. Nat Methods 16, 778–786 (2019).

13. Kerr, J.N.D. & Denk, W. Imaging in vivo: watching the brain in action. Nat Rev Neurosci 9, 195–205 (2008).

14. Ji, N., Freeman, J. & Smith, S.L. Technologies for imaging neural activity in large volumes. Nat Neurosci 19, 1154 (2016).

15. Yang, W. & Yuste, R. In vivo imaging of neural activity. Nat Methods 14, 349 (2017).

16. Weisenburger, S. & Vaziri, A. A Guide to Emerging Technologies for Large-Scale and Whole-Brain Optical Imaging of Neuronal Activity. Annual Review of Neuroscience 41, 431–452 (2018).

17. Chen, B.-C. et al. Lattice light-sheet microscopy: Imaging molecules to embryos at high spatiotemporal resolution. Science 346, 1257998 (2014).

18. Bouchard, M.B. et al. Swept confocally-aligned planar excitation (SCAPE) microscopy for high-speed volumetric imaging of behaving organisms. Nature Photonics 9, 113 (2015).

19. Voleti, V. et al. Real-time volumetric microscopy of in vivo dynamics and large-scale samples with SCAPE 2.0. Nat Methods 16, 1054–1062 (2019).

20. Ahrens, M.B., Orger, M.B., Robson, D.N., Li, J.M. & Keller, P.J. Whole-brain functional imaging at cellular resolution using light-sheet microscopy. Nat Methods 10, 413 (2013).

21. Tomer, R. et al. SPED Light Sheet Microscopy: Fast Mapping of Biological System Structure and Function. Cell 163, 1796–1806 (2015).

22. Holekamp, T.F., Turaga, D. & Holy, T.E. Fast Three-Dimensional Fluorescence Imaging of Activity in Neural Populations by Objective-Coupled Planar Illumination Microscopy. Neuron 57, 661–672 (2008).

23. Levoy, M., Ng, R., Adams, A., Footer, M. & Horowitz, M. Light Field Microscopy. ACM Transactions on Graphics 25, 924–934 (2006).

24. Broxton, M. et al. Wave optics theory and 3-D deconvolution for the light field microscope. Opt. Express 21, 25418–25439 (2013).

25. Prevedel, R. et al. Simultaneous whole-animal 3D imaging of neuronal activity using light-field microscopy. Nat Methods 11, 727–730 (2014).

26. Li, H. et al. Fast, volumetric live-cell imaging using high-resolution light-field microscopy. Biomed. Opt. Express 10, 29–49 (2019).

27. Pégard, N.C. et al. Compressive light-field microscopy for 3D neural activity recording. Optica 3, 517–524 (2016).

28. Nöbauer, T. et al. Video rate volumetric Ca2+ imaging across cortex using seeded iterative demixing (SID) microscopy. Nat Methods 14, 811 (2017).

29. Taylor, M.A., Nöbauer, T., Pernia-Andrade, A., Schlumm, F. & Vaziri, A. Brain-wide 3D light-field imaging of neuronal activity with speckle-enhanced resolution. Optica 5, 345–353 (2018).

30. Wagner, N. et al. Instantaneous isotropic volumetric imaging of fast biological processes. Nat Methods 16, 497–500 (2019).

31. Cong, L. et al. Rapid whole brain imaging of neural activity in freely behaving larval zebrafish (Danio rerio). eLife 6, e28158 (2017).

32. Hardy, J.W. Adaptive Optics for Astronomical Telescopes. (Oxford University Press, 1998).

33. Llavador, A., Sola-Pikabea, J., Saavedra, G., Javidi, B. & Martínez-Corral, M. Resolution improvements in integral microscopy with Fourier plane recording. Opt. Express 24, 20792–20798 (2016).

34. Guo, C., Liu, W., Hua, X., Li, H. & Jia, S. Fourier light-field microscopy. Opt. Express 27, 25573–25594 (2019).

35. Mertz, J. Optical sectioning microscopy with planar or structured illumination. Nat Methods 8, 811–819 (2011).

36. Nguyen, J.P. et al. Whole-brain calcium imaging with cellular resolution in freely behaving Caenorhabditis elegans. Proc Natl Acad Sci U S A 113, E1074–1081 (2016).

37. Venkatachalam, V. et al. Pan-neuronal imaging in roaming Caenorhabditis elegans. Proc Natl Acad Sci U S A 113, E1082–1088 (2016).

38. Kim, D.H. et al. Pan-neuronal calcium imaging with cellular resolution in freely swimming zebrafish. Nat Methods 14, 1107–1114 (2017).

39. Thouvenin, O. & Wyart, C. Tracking microscopy enables whole-brain imaging in freely moving zebrafish. Nat Methods 14, 1041 (2017).

40. Semmelhack, J.L. et al. A dedicated visual pathway for prey detection in larval zebrafish. Elife 3, e04878 (2014).

41. Henriques, P.M., Rahman, N., Jackson, S.E. & Bianco, I.H. Nucleus isthmi is required to sustain target pursuit during visually guided prey-catching. Current Biology 29, 1771–1786 (2019).

42. Bianco, I.H. & Engert, F. Visuomotor transformations underlying hunting behavior in zebrafish. Current biology 25, 831–846 (2015).

43. Muto, A. et al. Activation of the hypothalamic feeding centre upon visual prey detection. Nature communications 8, 15029 (2017).

44. Muto, A., Ohkura, M., Abe, G., Nakai, J. & Kawakami, K. Real-Time Visualization of Neuronal Activity during Perception. Current Biology 23, 307–311 (2013).

45. Kim, D.H. et al. Pan-neuronal calcium imaging with cellular resolution in freely swimming zebrafish. Nat Methods 14, 1107 (2017).

46. Zhou, P. et al. Efficient and accurate extraction of in vivo calcium signals from microendoscopic video data. eLife 7, e28728 (2018).

47. Pnevmatikakis, Eftychios A., et al. Simultaneous Denoising, Deconvolution, and Demixing of Calcium Imaging Data. Neuron 89, 285–299 (2016).

48. Mickoleit, M. et al. High-resolution reconstruction of the beating zebrafish heart. Nat Methods 11, 919 (2014).

49. Grutzendler, J. & Nedergaard, M. Cellular Control of Brain Capillary Blood Flow: In Vivo Imaging Veritas. Trends in Neurosciences 42, 528–536 (2019).

50. Kornfield, T.E. & Newman, E.A. Measurement of retinal blood flow using fluorescently labeled red blood cells. Eneuro 2 (2015).

51. Dunsby, C. Optically sectioned imaging by oblique plane microscopy. Opt. Express 16, 20306–20316 (2008).

52. Iadecola, C. The Neurovascular Unit Coming of Age: A Journey through Neurovascular Coupling in Health and Disease. Neuron 96, 17–42 (2017).

53. Zhang, B., Yao, Y., Zhang, H., Kawakami, K. & Du, J. Left Habenula Mediates Light-Preference Behavior in Zebrafish via an Asymmetrical Visual Pathway. Neuron 93, 914–928 (2017).

54. Filosa, A., Barker, A.J., Dal Maschio, M. & Baier, H. Feeding State Modulates Behavioral Choice and Processing of Prey Stimuli in the Zebrafish Tectum. Neuron 90, 596–608 (2016).

55. Khoobehi, B., Peyman, G.A., Carnahan, L.G. & Hayes, R.L. A Novel Approach for Freeze-Frame Video Determination of Volumetric Blood Flow in the Rat Retina. Ophthalmic Surgery Lasers & Imaging 34, 505–514 (2003).

56. Smith, S.L. & Häusser, M. Parallel processing of visual space by neighboring neurons in mouse visual cortex. Nat Neurosci 13, 1144–1149 (2010).

## References

52. Zhang, B., Yao, Y., Zhang, H., Kawakami, K. & Du, J. Left Habenula Mediates Light-Preference Behavior in Zebrafish via an Asymmetrical Visual Pathway. Neuron 93, 914–928 (2017).

53. Filosa, A., Barker, A.J., Dal Maschio, M. & Baier, H. Feeding State Modulates Behavioral Choice and Processing of Prey Stimuli in the Zebrafish Tectum. Neuron 90, 596–608 (2016).

54. Khoobehi, B., Peyman, G.A., Carnahan, L.G. & Hayes, R.L. A Novel Approach for Freeze-Frame Video Determination of Volumetric Blood Flow in the Rat Retina. Ophthalmic Surgery Lasers & Imaging 34, 505–514 (2003).

55. Smith, S.L. & Häusser, M. Parallel processing of visual space by neighboring neurons in mouse visual cortex. Nat Neurosci 13, 1144–1149 (2010).

