## supplementary materials for "Capturing volumetric dynamics at high speed in the brain by confocal light field microscopy"


| | |
|---|---|
| Supplementary Figure 1 | Experimental implementation of Confocal LFM and 3D tracking system for larval zebrafish brain imaging |
| Supplementary Figure 2 | System synchronization |
| Supplementary Figure 3 | Design of the confocal detection mask |
| Supplementary Figure 4 | Experimentally measured PSFs in non-confocal and Confocal LFMs for zebrafish imaging |
| Supplementary Figure 5 | Experimentally measured 5 PSFs focusing at different depths and covering an extended axial range |
| Supplementary Figure 6 | Characterization of resolution afforded by outermost micro-lens in Confocal LFM for zebrafish imaging |
| Supplementary Figure 7 | Comparison of reconstructions with and without system correction |
| Supplementary Figure 8 | Characterization of field distortion |
| Supplementary Figure 9 | Characterization of reconstruction in Confocal LFM for zebrafish imaging |
| Supplementary Figure 10 | Comparison of raw images of fluorescent beads captured in non-confocal LFM and Confocal LFM |
| Supplementary Figure 11 | Characterization of calcium activities on activated neurons and brain regions in larval zebrafish during its prey capture behavior |

| | |
|---|---|
| Supplementary Figure 12 | Schematics of Confocal LFM optimized for mice brain imaging |
| Supplementary Figure 13 | Experimentally measured PSFs in non-confocal and Confocal LFMs for mice imaging |
| Supplementary Figure 14 | Characterization of resolution afforded by outermost micro-lens in Confocal LFM for mice imaging |
| Supplementary Figure 15 | Characterization of reconstruction in Confocal LFM optimized for mice brain imaging |
| Supplementary Figure 16 | Comparison of Confocal LFM and non-confocal LFM imaging in awake mice brain |
| Supplementary Figure 17 | Extraction of activity traces and spatial footprints of neural structures by CNMF-E |
| Supplementary Figure 18 | Characterization of photobleaching in functional imaging in awake mice brain using Confocal LFM |
| Supplementary Table 1 | Acquisition parameters for Confocal LFM imaging in zebrafish |
| Supplementary Table 2 | Acquisition parameters for Confocal LFM imaging in mice |
| Supplementary Table 3 | Acquisition parameters for two photon imaging in mice |
| Supplementary Video 1 | Animated schematics of Confocal LFM with fast axial scanning |
| Supplementary Video 2 | Comparison of Confocal LFM and non-confocal LFM when imaging spontaneous neural activities in restrained larval zebrafish brain |
| Supplementary Video 3 | 3D volumetric imaging of larval zebrafish brain |
| Supplementary Video 4 | Whole brain functional imaging in larval zebrafish under light stimulation |
| Supplementary Video 5 | Whole brain functional imaging of a freely swimming larval zebrafish during prey capture behavior |
| Supplementary Video 6 | Unbiased volumetric reconstruction of active neural structures in awake mouse brain imaged by Confocal LFM |
| Supplementary Video 7 | Active neural structures extracted by CNMF-E on each z plane independently in an imaging volume captured by Confocal LFM |

| | |
|---|---|
| Supplementary Video 8 | Imaging circulating blood cells in awake mouse brain by Confocal LFM |
| Supplementary Video 9 | Imaging circulating blood cells in awake mouse brain over extended period of time |
| Supplementary Video 10 | Analysis of instantaneous speeds of circulating cells in blood vessels |

**Supplementary Figure 1| Experimental implementation of Confocal LFM and 3D tracking system for larval zebrafish brain imaging**

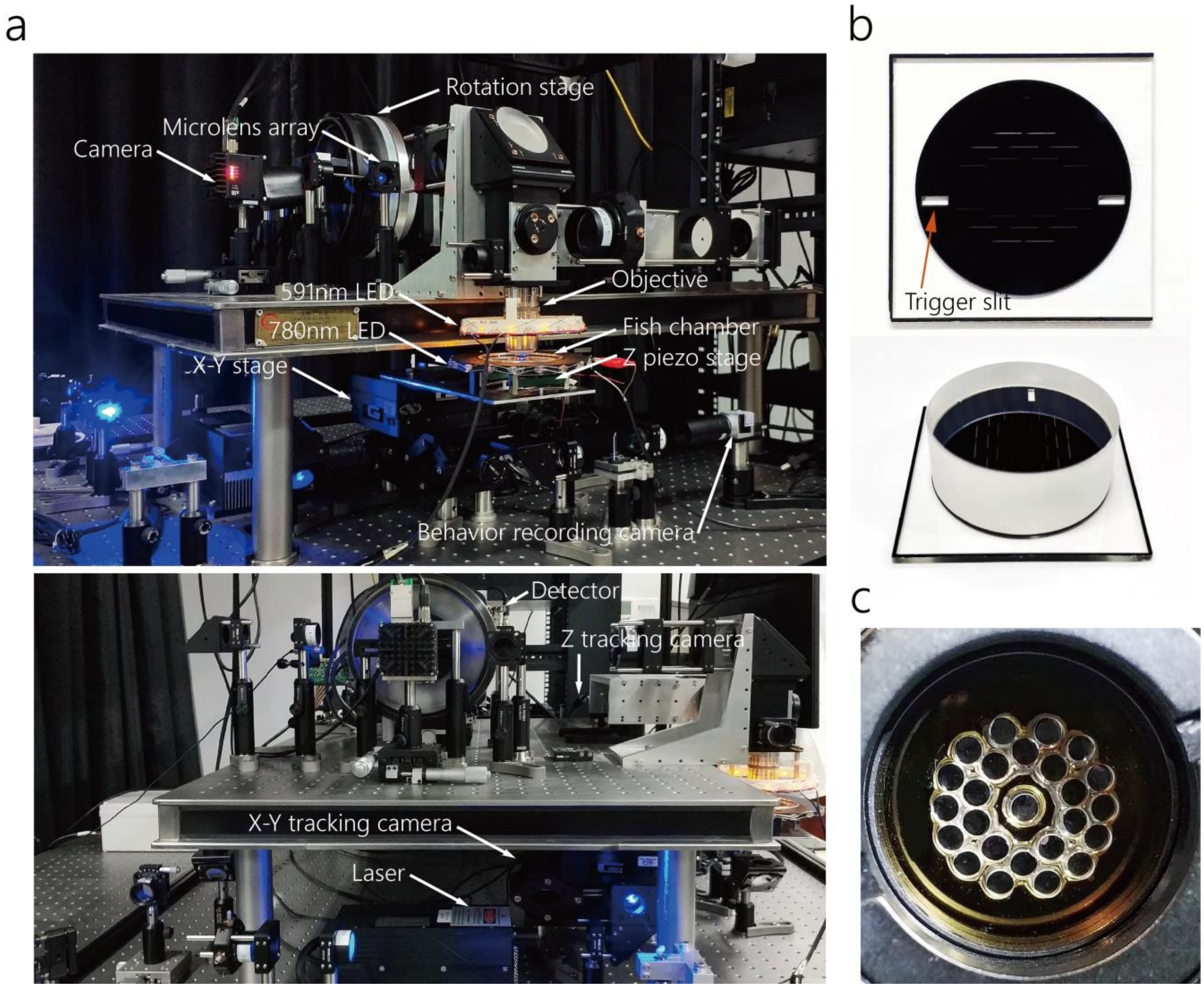


(a) Overviews of the Confocal LFM and 3D tracking system. (b) Top and side views of a representing mask with glass plate attached. (c) Micro-lens array consisting of 25 micro-lenses housed in a customized aluminum mount.

**Supplementary Figure 2| System synchronization**

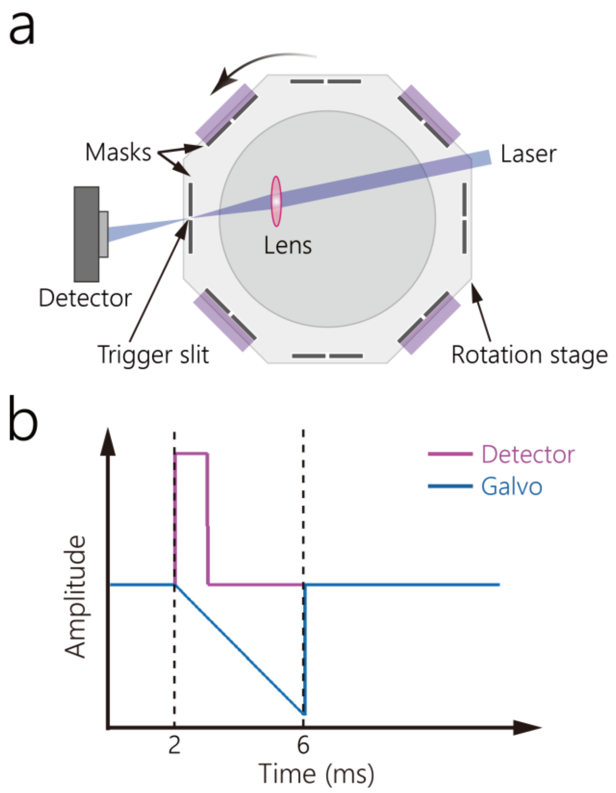


(a) Schematics of trigger generation for system synchronization. A separate 450nm laser was focused on to the mask. (b) When the trigger slit (Supplementary Figure 1b) on the mask was rotated to this focused laser spot, a detector sitting on the other side of the mask generated a step rising signal upon the transmission of the focused laser beam to serve as the trigger to synchronize scanning galvo, laser power and camera exposure. When camera ran at 30 Hz frame rate, the scanning and exposure time (between dashed lines) was ~4ms.

**Supplementary Figure 3| Design of the confocal detection mask**

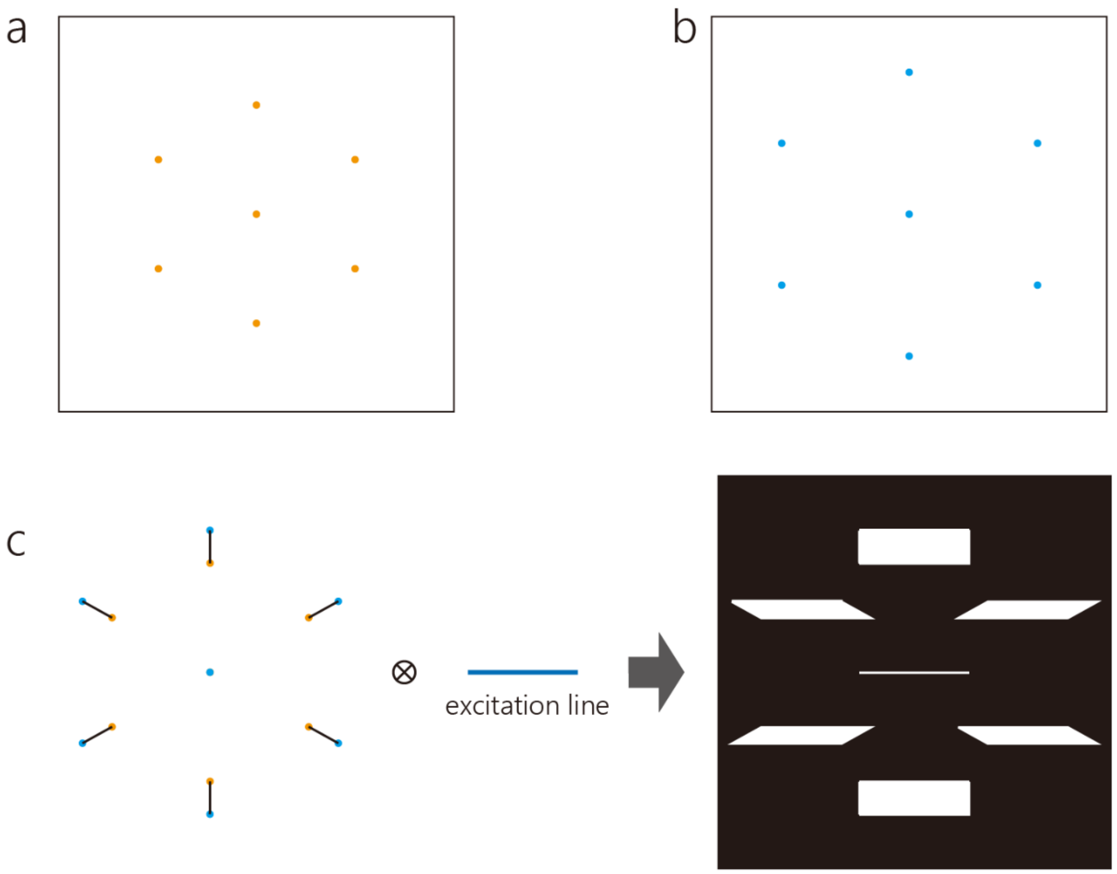


(a) The image of a fluorescent bead formed by the micro-lens array. The bead was placed in the center of the field of view in x-y directions and at the top axial edge $z_{top}$ of the designed imaging volume. (b) The image of the same bead as in (a) when it was placed at the bottom axial edge $z_{bottom}$ of the designed imaging volume. (c) Draw lines between corresponding bead images in (a) and (b) to form a new image, and convolve this new image with a line, representing the excitation laser beam focused by a cylindrical lens, to generate the pattern of confocal detection mask. The black region is opaque and blocks background fluorescence from out-of-focus volume.

**Supplementary Figure 4| Experimentally measured PSFs in non-confocal and Confocal LFMs for zebrafish imaging**

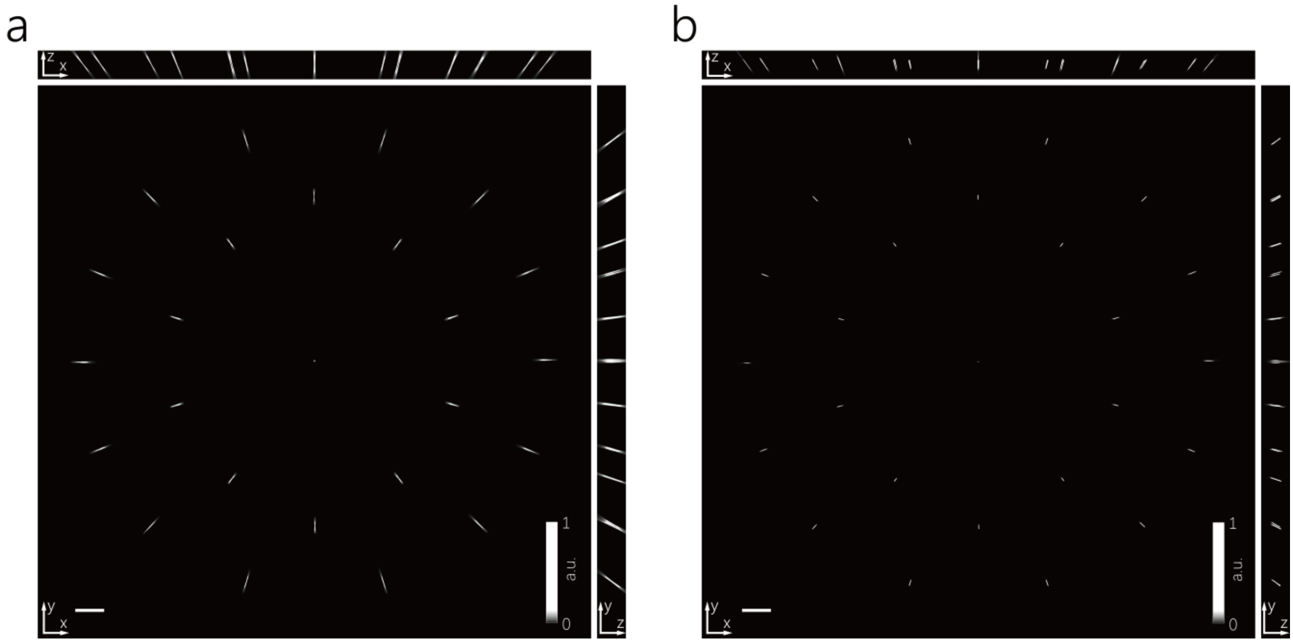


Maximum Intensity Projections (MIPs) of PSFs measured in non-confocal LFM (a, confocal detection mask was removed) and Confocal LFM (b, confocal detection mask was present). The PSFs were imaged as 3D stacks on a fluorescent bead of 1 µm diameter when it was scanned in z direction. Scale bar, 200 µm.

**Supplementary Figure 5| Experimentally measured 5 PSFs focusing at different depths and covering an extended axial range**

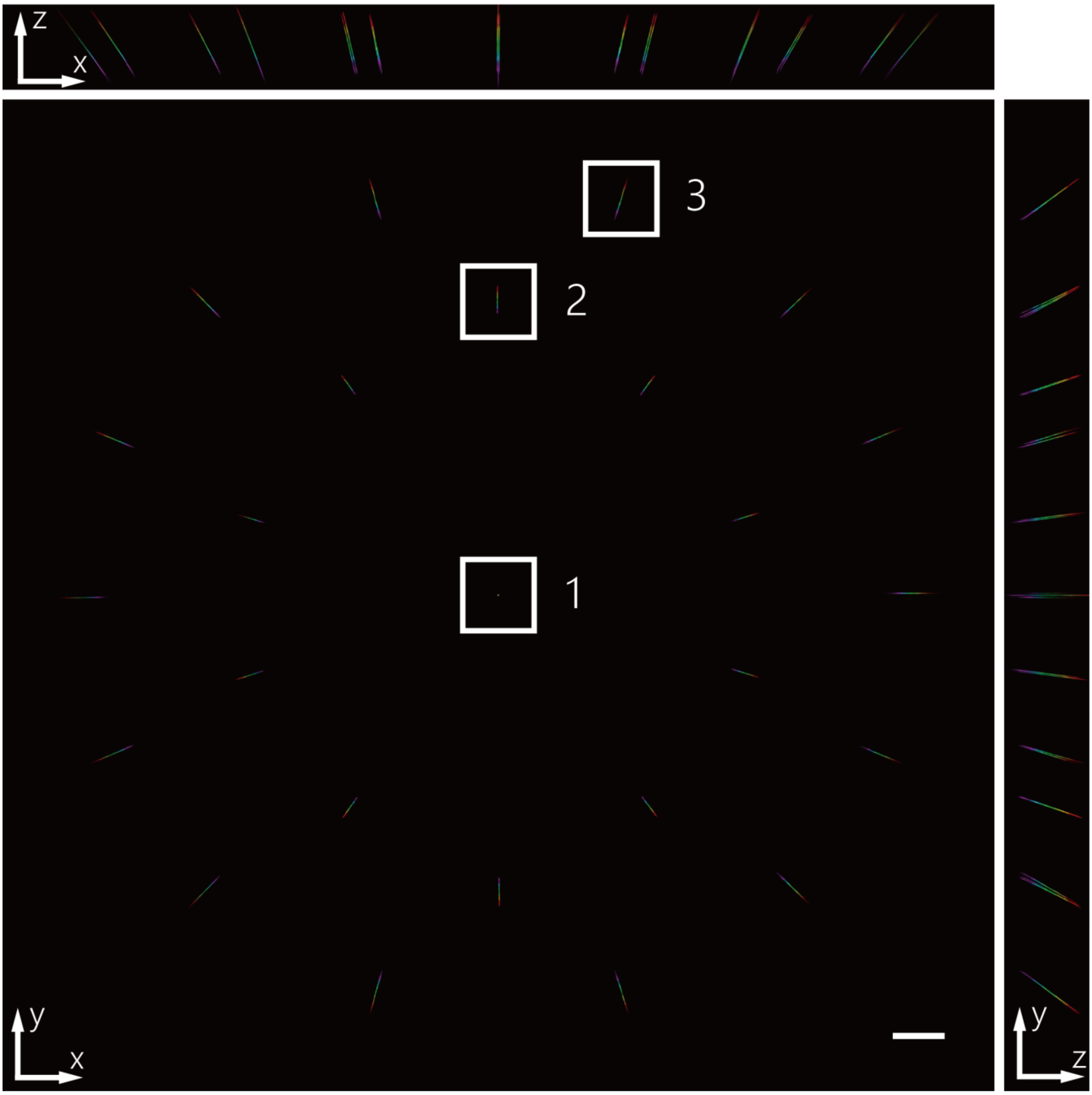


MIPs of 5 PSFs measured from 5 different mask-glass plate combinations that were designed to focus at different depths and, when used together, can continuously cover an extended axial range. These 5 PSFs were colored in red, yellow, green, cyan and magenta, respectively. They were merged for display. The magnified views of region 1, 2 and 3 were shown in Figure 1b in the main text. Scale bar, 200 μm.

**Supplementary Figure 6| Characterization of resolution afforded by outermost micro-lens in Confocal LFM for zebrafish imaging**

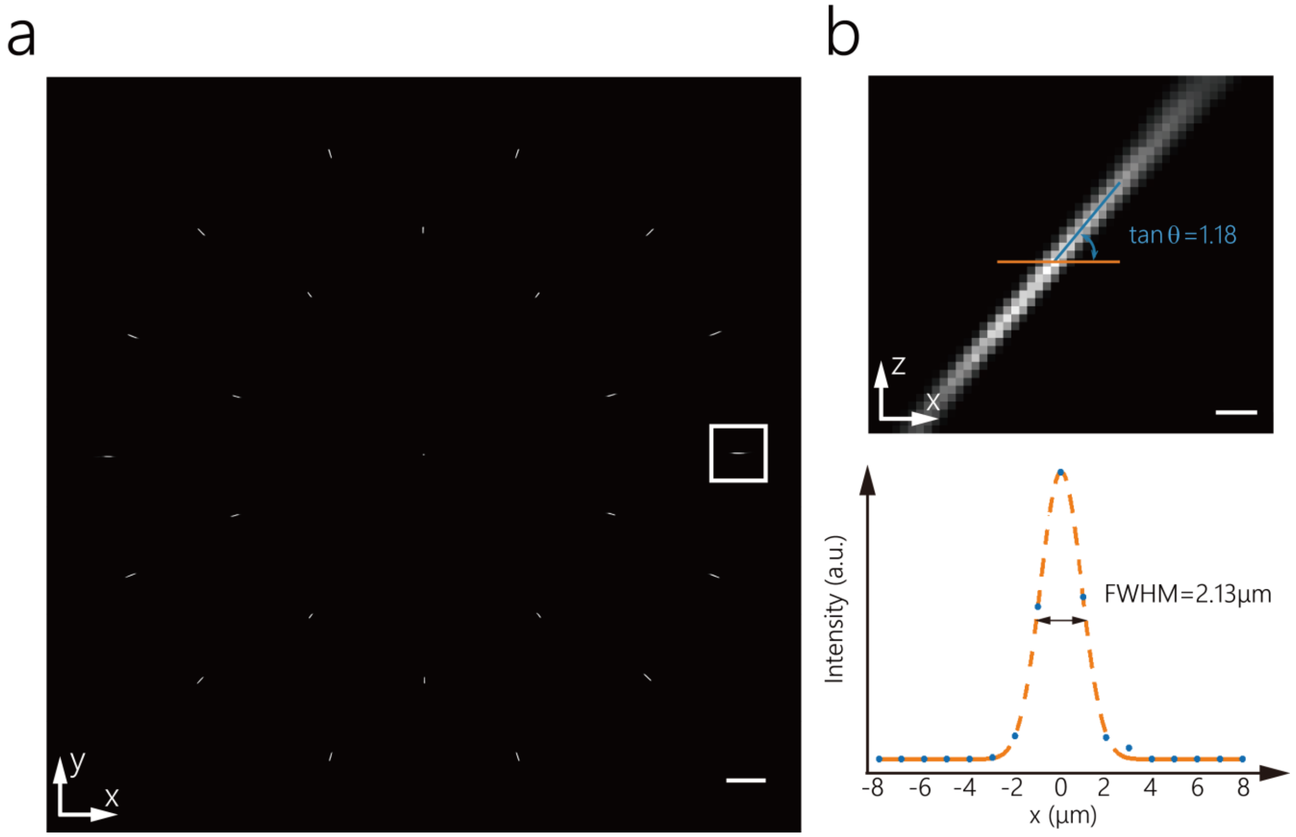


(a) MIP of the PSF measured from one example mask. Scale bar, 200 μm. (b) Zoom-in side view of the PSF measured from a representative outermost micro-lens, as indicated by the box in (a). Scale bar, 10 μm. The line profile (orange line) of the PSF waist was fitted by Gaussian function (bottom) and indicated a measured Full Width at Half Maximum (FWHM) of 2.13 μm, which was close to theoretical estimation of 2 μm. The tilt angle of the PSF in z direction was found to be $\theta \approx 50°$. The axial resolution afforded by this micro-lens could be estimated as $2.13 \times \tan\theta \approx 2.5$ μm, because two beads with 2.5 μm inter spacing in z direction would be laterally separated by 2.13 μm in the image formed by this outermost micro-lens, which is within the lateral resolution limit. When combining information from all micro-lens correctly, the whole system could achieve this optimal resolution afforded by this micro-lens.

**Supplementary Figure 7| Comparison of reconstructions with and without system correction**

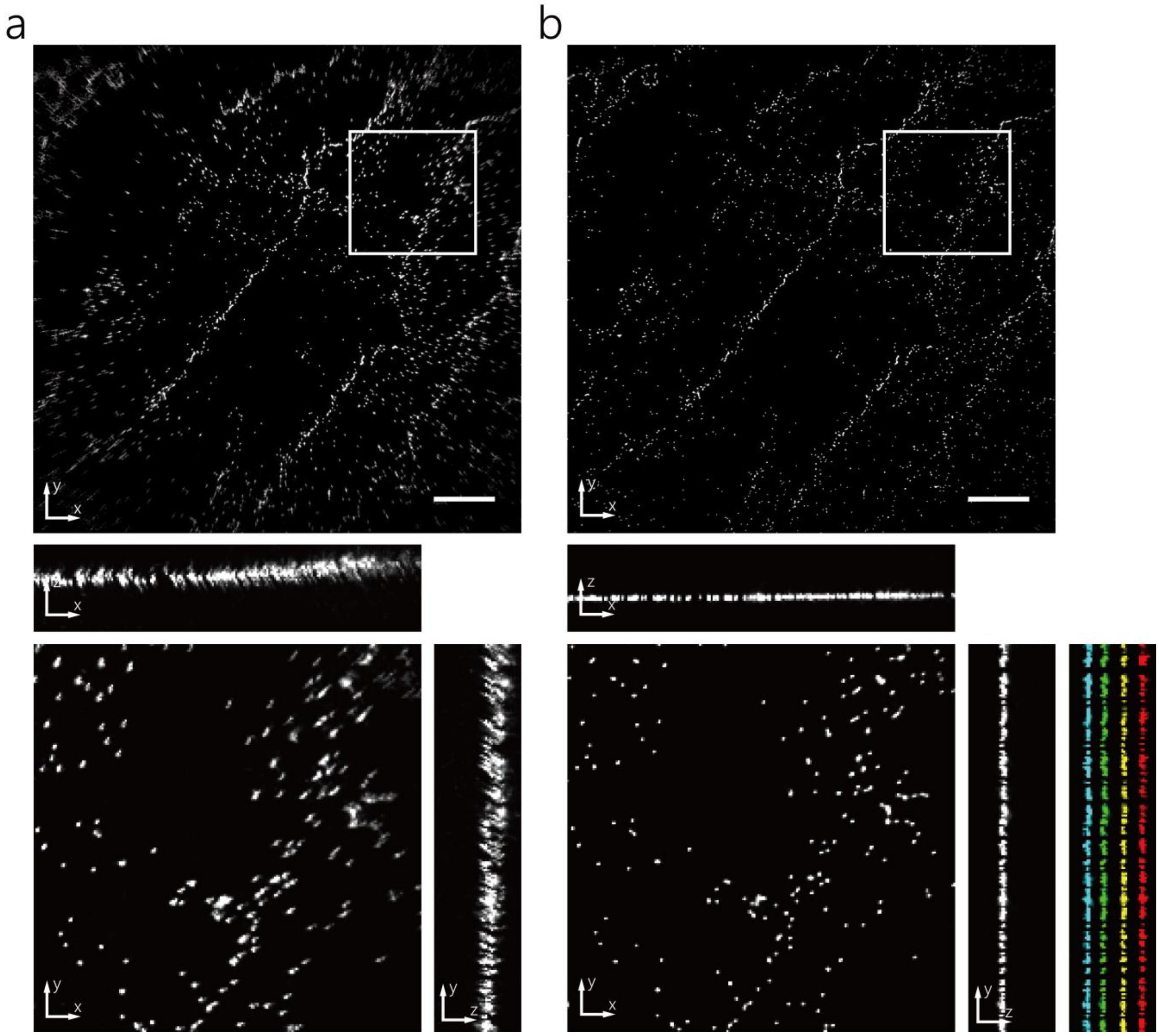


Overviews (top) and zoom-in views (bottom) of reconstructed images of densely packed 1 µm diameter fluorescent beads on a glass slide without (a) and with (b) system correction. (a) Without system correction, the reconstruction was not optimal, as evidenced by the elongated shape of reconstructed beads. (b) After applying system correction, reconstruction quality was improved and small spatial footprints of individual beads were more correctly restored. This correction worked well throughout the entire imaging volume. Four reconstructions were carried out on the beads slide placed at different z positions with 10 µm interspacing and colored in cyan, green, yellow & red. Scale bar, 100 µm.

**Supplementary Figure 8| Characterization of field distortion**

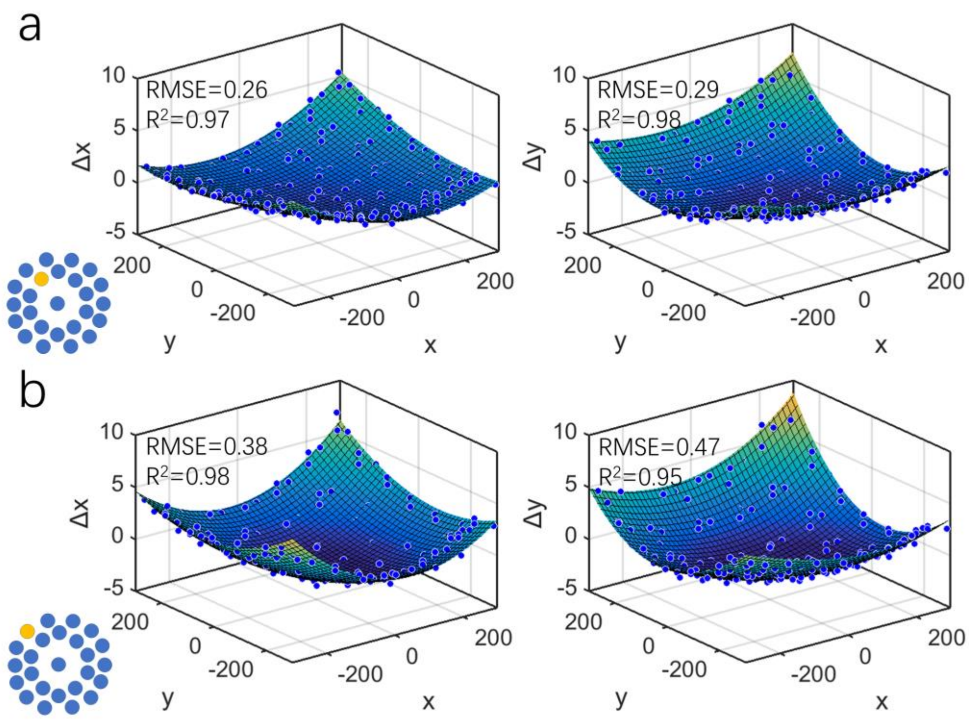


Experimental measuring and fitting of field distortions on two example micro-lenses. The selected micro-lenses were indicated in yellow in the lower-right corner insets in (a) and (b). To measure the field distortion, a layer of sparsely spread 1 µm diameter fluorescent beads near the focal plane were imaged. The images of these beads measured from different peripheral micro-lenses were compared with that obtained from the central micro-lens. Due to the field distortion, these images could not match exactly. The relative positional shifts of these beads (individual blue spots) in x and y directions (Δx and Δy) could be characterized, as shown above. These shifts varied depending on their locations and could be well fitted with quadratic polynomial surfaces.

**Supplementary Figure 9| Characterization of reconstruction in Confocal LFM for zebrafish imaging**

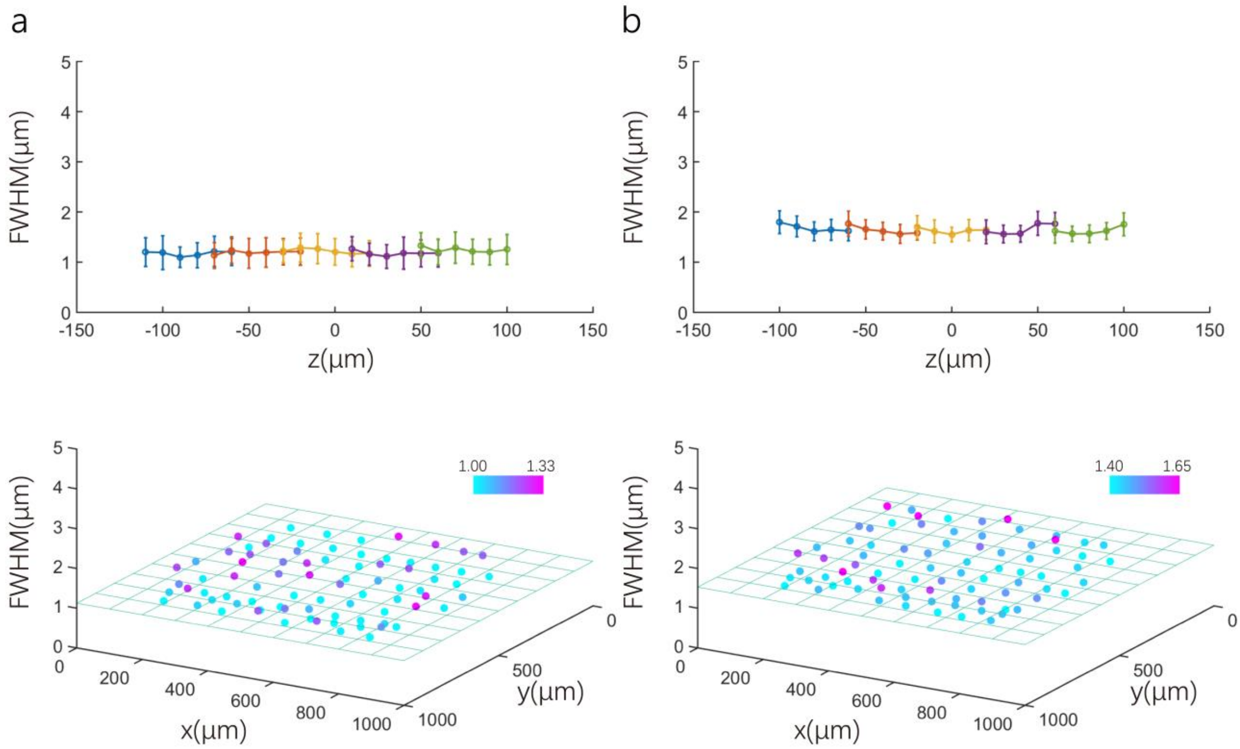


The performance of the reconstruction was characterized by imaging sparsely spread 0.5 μm diameter fluorescent beads on a glass slide placed at different z depths. With system correction, reconstructions of these isolated beads resulted in spot like images with FWHM of ~1.2 μm in lateral direction (a) and ~1.6 μm in axial direction (b) across the whole imaging volume of ⌀ 800 μm x 200 μm. Top, FWHM of x-profile (a) and z-profile (b) of reconstructed beads across 200 μm in axial direction. Bottom, FWHM of x-profile (a) and z-profile (b) of reconstructed beads across ⌀ 800 μm in x-y plane near focal plane. Because this size was smaller than the diffraction limit of 2.13 x 2.13 x 2.5 $\mu m^3$, the reconstruction was optimal and diffraction limited.

**Supplementary Figure 10| Comparison of raw images of fluorescent beads captured in non-confocal LFM and Confocal LFM**

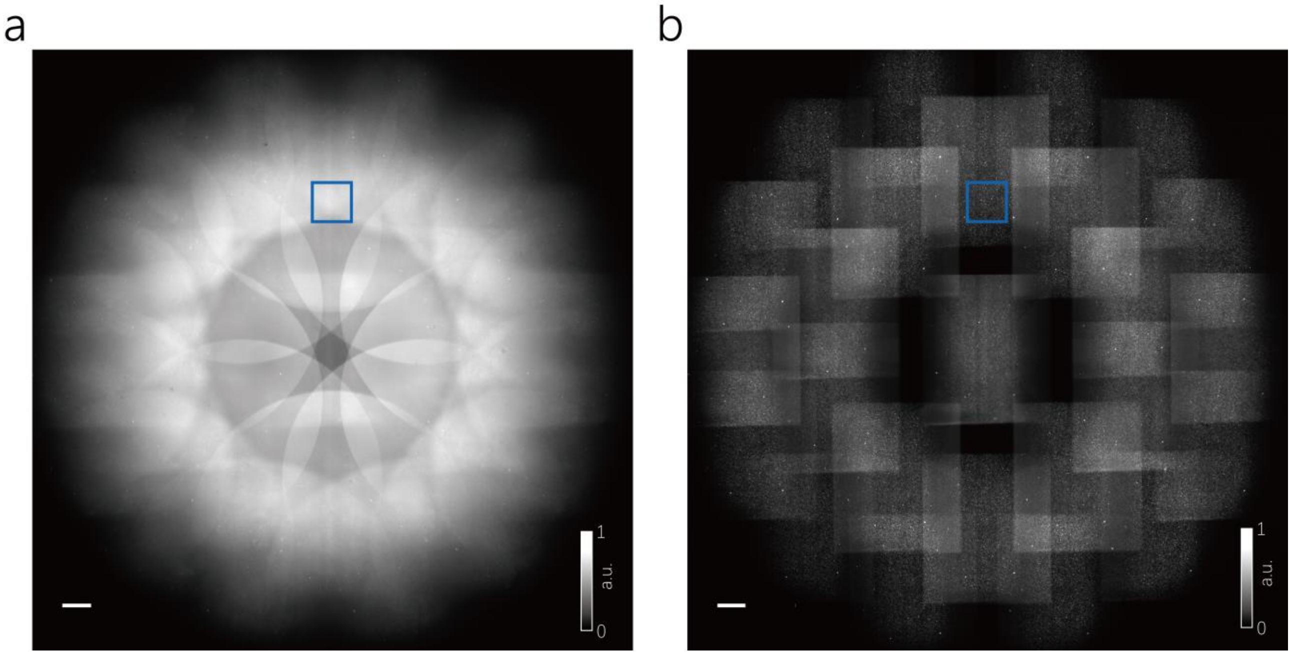


Raw images captured on the camera in non-confocal LFM (a, confocal detection mask was removed) and Confocal LFM (b, confocal detection mask was present). The fluorescent beads are 0.5μm in diameter and randomly distributed in 3D agarose gel of about 1 mm thick. The magnified images of indicated regions in blue boxes were shown in Figure 1c in the main text. Scale bar, 200 μm.

**Supplementary Figure 11| Characterization of calcium activities on activated neurons and brain regions in larval zebrafish during its prey capture behavior**

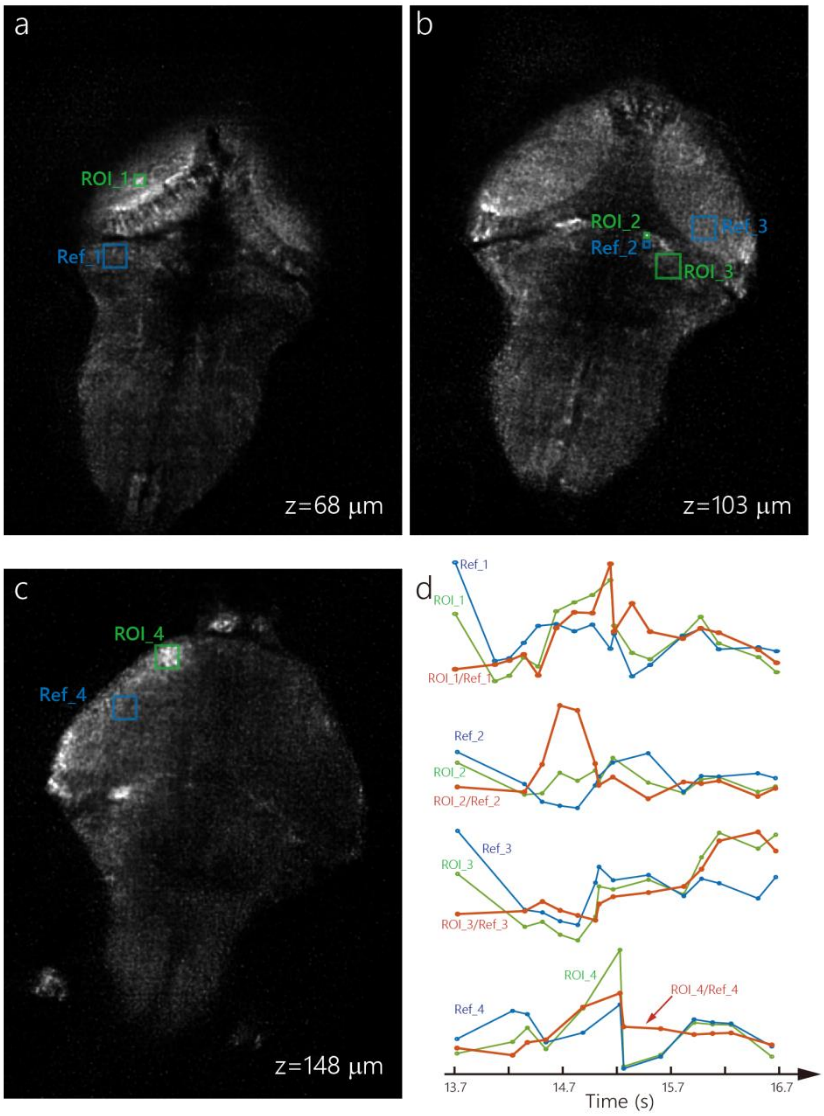


To normalize the imaging intensity change caused by body rotations during larval zebrafish's swimming, the calcium activities on activated neurons and brain regions were inferred as ratios of fluorescent intensities measured from the regions of interests (ROI_1, 2, 3 & 4, indicated by green boxes in (a), (b) and (c)) and over the respective reference regions (Ref_1, 2, 3 & 4, indicated by blue boxes in (a), (b) and (c)). The reference regions were selected manually based on two criteria: they were relatively close to the regions of interests and they had no obvious neural activation as identified by patterned fluorescent signal change relative to their surroundings. (d) Raw traces of fluorescent intensity change in regions of interests (green), their corresponding reference regions (blue) and the ratios between them (red), which were the same as that in Figure 3f in the main text.

**Supplementary Figure 12| Schematics of Confocal LFM optimized for mice brain imaging**

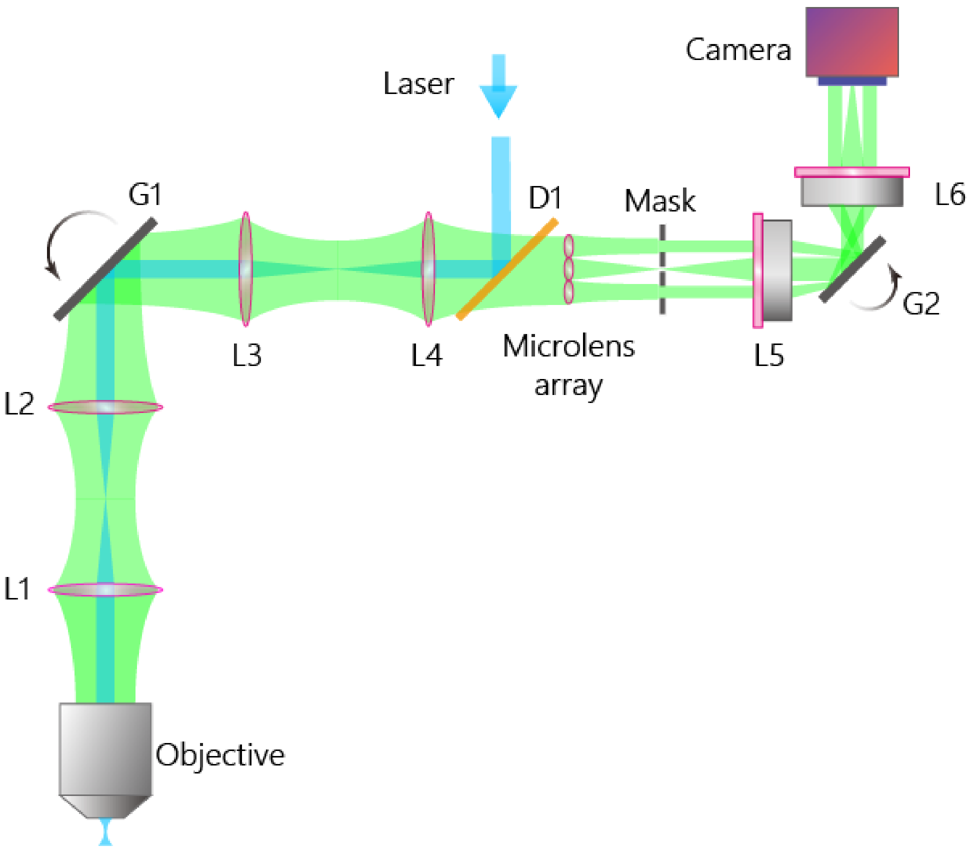


L1-L4: achromatic optical relay lenses; D1: long pass dichroic mirror; L5, L6 telecentric imaging lenses; G1, G2 scanning galvo mirrors. Detailed description of system design and parameters of all components could be found in Methods.

**Supplementary Figure 13| Experimentally measured PSFs in non-confocal and Confocal LFMs for mice imaging**

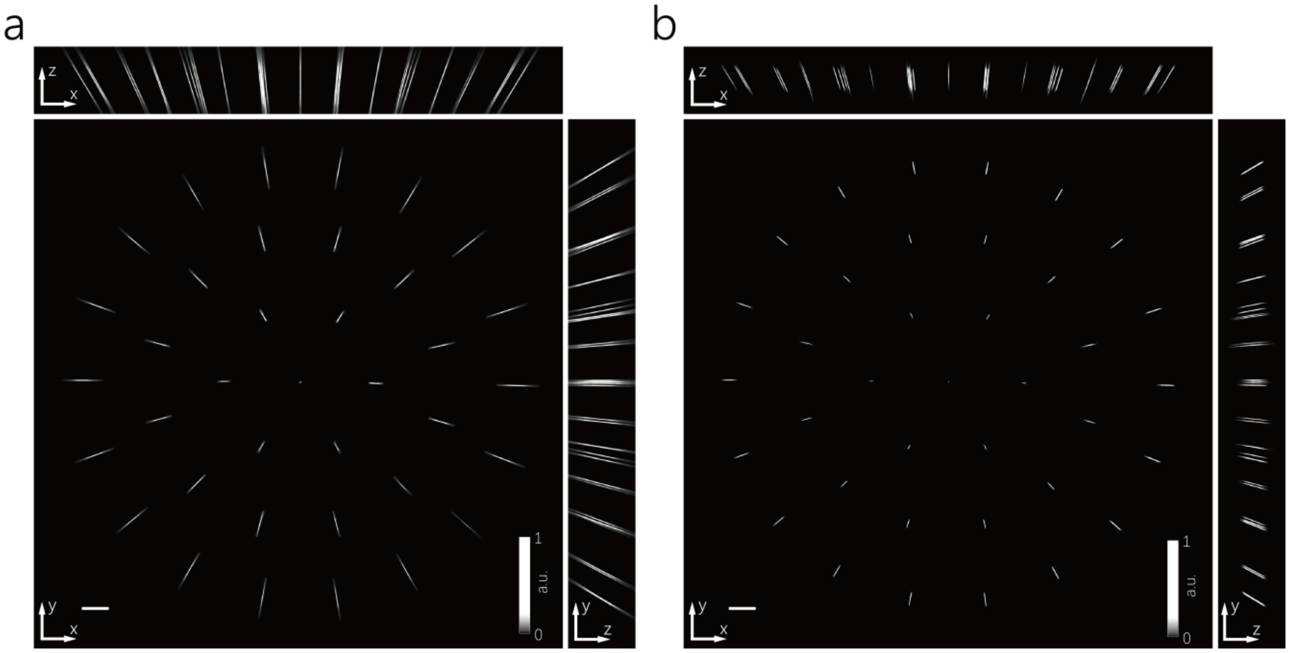


Maximum Intensity Projections (MIPs) of PSFs measured in non-confocal LFM (a, confocal detection mask was removed) and Confocal LFM (b, confocal detection mask was present). The PSFs were imaged as 3D stacks on a fluorescent bead of 1 µm diameter when it was scanned in z direction. Scale bar, 200 µm.

**Supplementary Figure 14| Characterization of resolution afforded by outermost micro-lens in Confocal LFM for mice imaging**

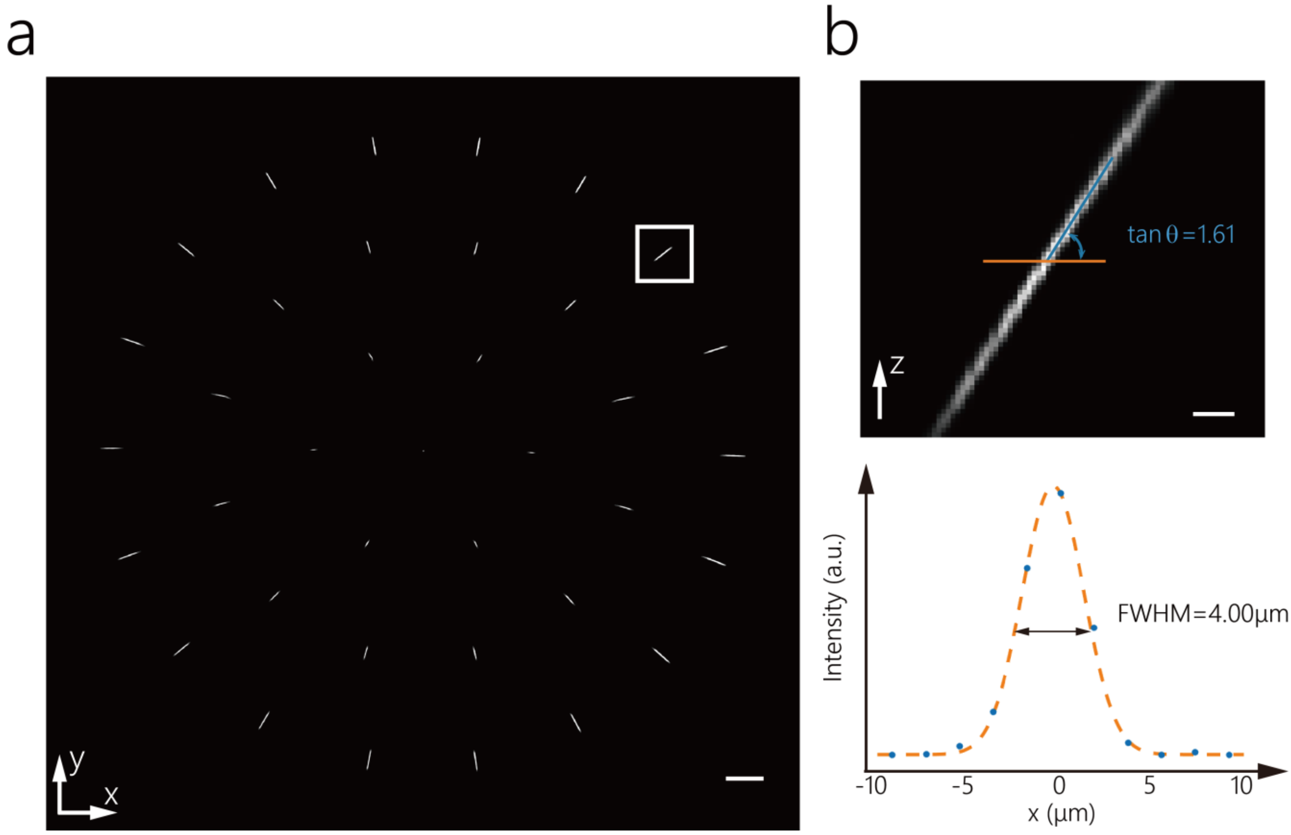


(a) MIP of the PSF. Scale bar, 200 μm. (b) Zoom-in side view of the PSF measured from a representative outermost micro-lens, as indicated by the box in (a). Scale bar, 20 μm. The line profile (orange line) of the PSF waist was fitted by Gaussian function (bottom) and indicated a measured FWHM of 4.0 μm, which was close to theoretical estimation of 3.8 μm. The tilt angle of the PSF in z direction was found to be $\theta \approx 58°$. The axial resolution afforded by this micro-lens could be estimated as $4 \times \tan\theta \approx 6.4\ \mu m$. When combining information from all micro-lens correctly, the whole system could achieve this optimal resolution afforded by this micro-lens.

**Supplementary Figure 15| Characterization of reconstruction in Confocal LFM optimized for mice brain imaging**

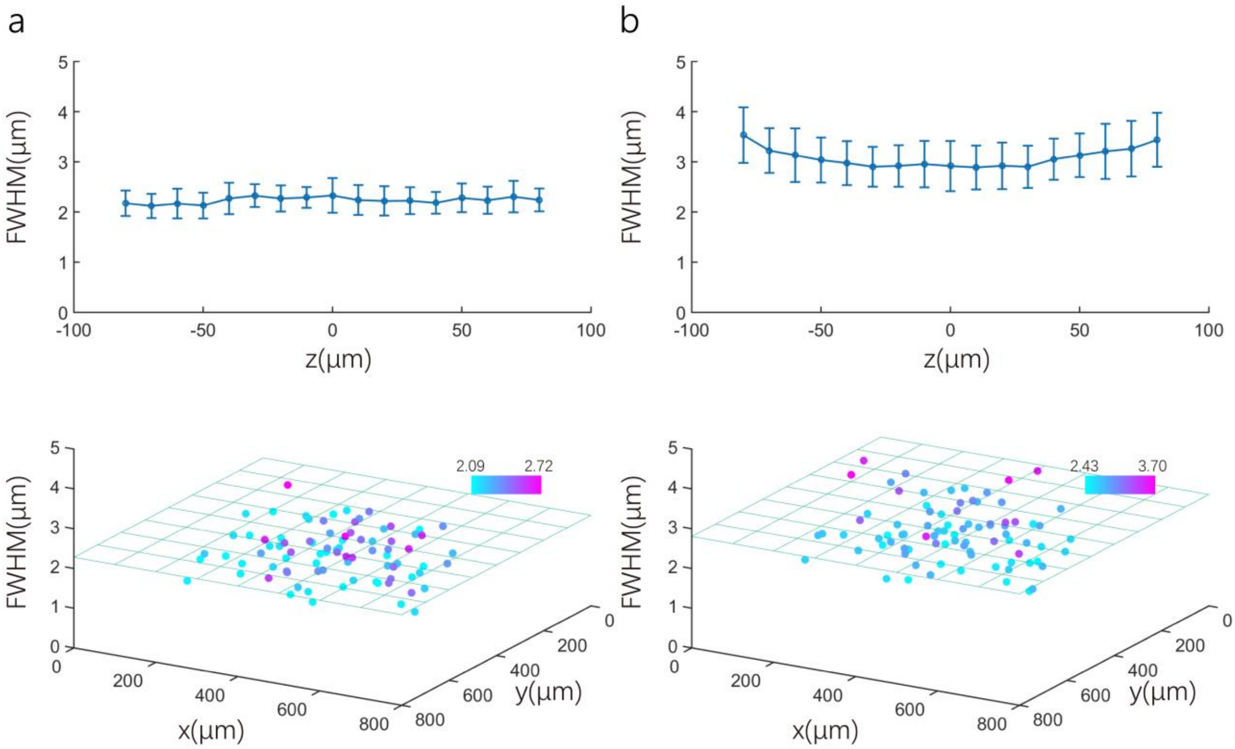


The performance of the reconstruction was characterized by imaging sparsely spread 1 μm diameter fluorescent beads on a glass slide placed at different z depths. With system correction, reconstructions of these isolated beads resulted in spot like images with FWHM of ~2.2 μm in lateral direction (a) and ~3.1 μm in axial direction (b) across the whole imaging volume of Φ 800 μm x 150 μm. Top, FWHM of x-profile (a) and z-profile (b) across 150 μm on z axis. Bottom, FWHM of x-profile (a) and z-profile (b) across Φ 800 μm on x-y plane near focal plane. Because this size was smaller than the diffraction limit of 3.8 x 3.8 x 6.0 $\mu m^3$, the reconstruction was optimal and diffraction limited.

**Supplementary Figure 16| Comparison of Confocal LFM and non-confocal LFM imaging in awake mice brain**

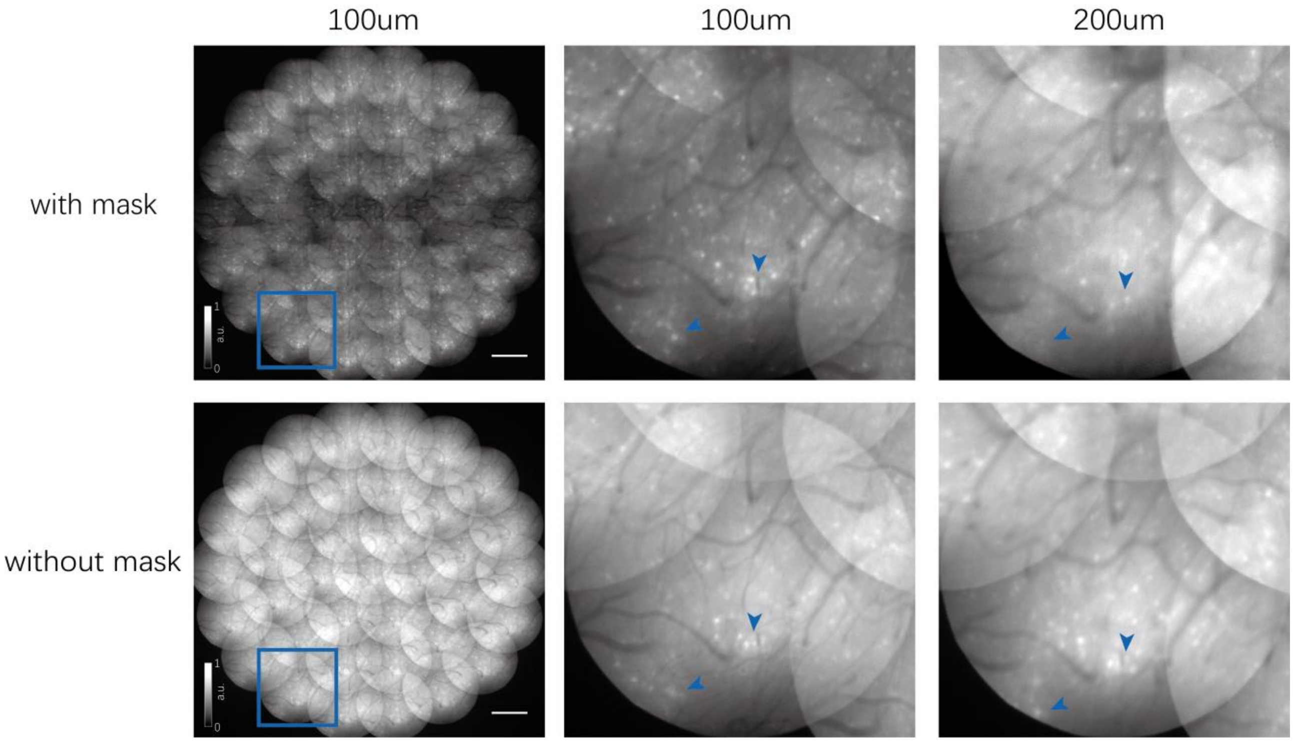


Example raw images captured by Confocal LFM (top, confocal detection mask was present) and non-confocal LFM (bottom, confocal detection mask was removed) in awake mouse brain when focusing at 100 µm below the cortex surface (left). Neurons in mouse brain was labeled with GCamp6s by virus injection. Zoom-in views of a small region indicated by blue boxes when focusing at 100 µm (middle) and 200 µm (right) below the cortex surface. Bright neurons (indicated by the arrow heads) near the surface could be well rejected when imaging at depth of 200 µm in Confocal LFM, however, they mixed together with in-focus neurons and cause interfering background in non-confocal LFM. Meanwhile, raw images captured by Confocal LFM had much suppressed background than non-confocal LFM. Scale bar, 500 µm.

**Supplementary Figure 17| Extraction of activity traces and spatial footprints of neural structures by CNMF-E**

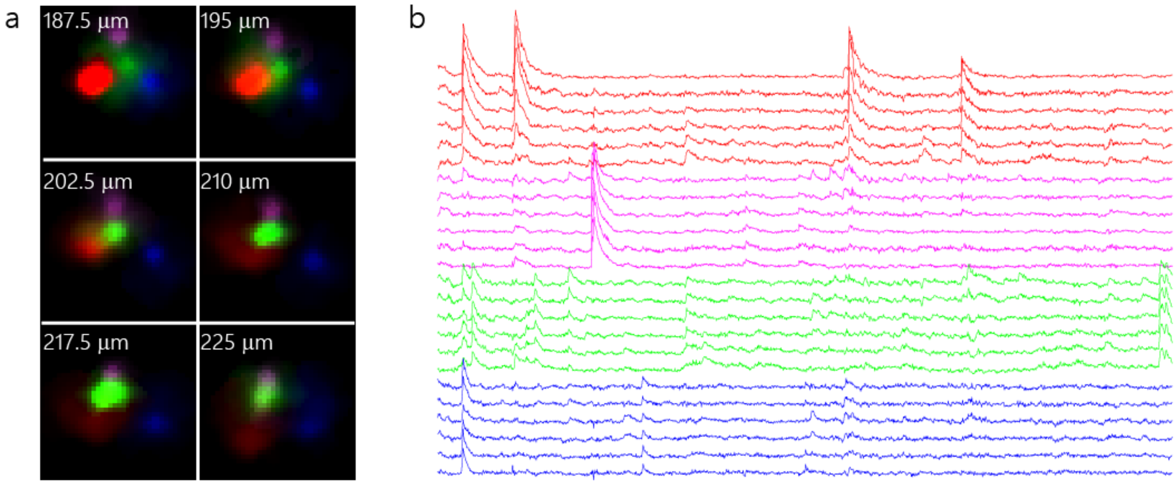


The neural activities and spatial footprints of densely packed neural structures captured by Confocal LFM in awake mouse brain were extracted by CNMF-E on 6 consecutive z planes independently, as shown in Figure 4d and 4e in the main text. The footprints of the same neural structure (a) could be identified by their highly correlated activity traces (b). 4 different neural structures were identified and their footprints and activity traces were indicated by 4 different colors.

**Supplementary Figure 18| Characterization of photobleaching in functional imaging in awake mice brain using Confocal LFM**

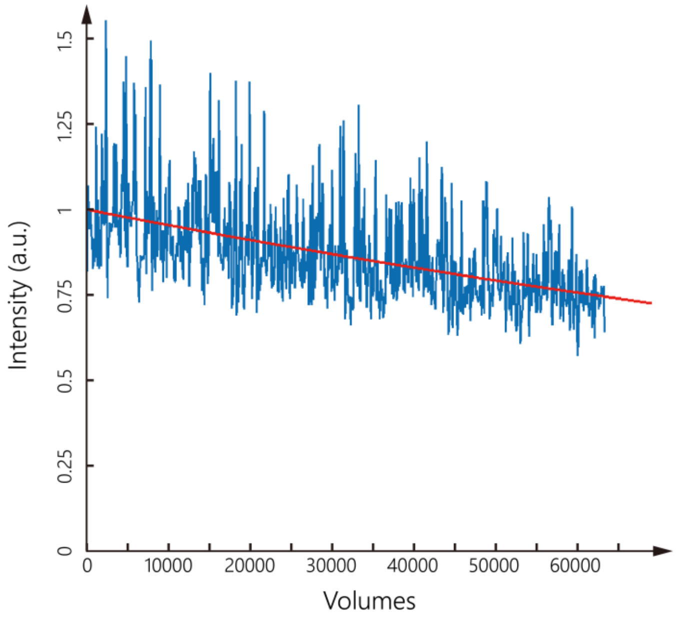


The intensity of fluorescence from GCamp6s labeled neurons in awake mouse brain decreased slowly when imaged by Confocal LFM continuously over 64,000 volumes. The illumination intensity of the excitation laser was about 0.73 $mW/mm^2$ (Supplementary Table 2) and Confocal LFM recorded neural activities at a speed of 5 volumes/s continuously over 4 hours (64000 volumes in total). The spontaneous activities of neurons in the mouse led to fluctuations in measured fluorescence intensity (blue, intensities were sampled at 640 time points in 64,000 volumes). Exponential fit (red) of this fluctuating curve showed that the fluorescence intensity decreased by 26% after 64,000 imaging volumes and the estimated decay constant was about 208,000 time points.

**Supplementary Table 1| Acquisition parameters for Confocal LFM imaging in zebrafish**

| Experiments | Experimental Models: Organisms/Strains | Age (dpf) | Volum e rate (Hz) | Camera frame Rate (Hz) | Exposure time (ms) | Average laser illumination intensity (mW/mm$^2$) |
|---|---|---|---|---|---|---|
| Figure 2a Supplementary Video 2 | *Tg(HuC:GCaMP6s)* | 6 | 1 | 2 | 8 | 1.2 |
| Figure 2b, c & d Supplementary Video 3, 4 | *Tg(HuC:GCaMP6s)* | 5 | 6 | 30 | 4 | 8.7 |
| Figure 3, Supplementary Figure 11 Supplementary Video 5 | *Tg(HuC:GCaMP6s)* | 8 | 6 | 30 | 4 | 8.7 |

**Supplementary Table 2| Acquisition parameters for Confocal LFM imaging in mice**

| Experiments | Mice strains/virus injection system | Age (weeks) | Volume rate (Hz) | Exposure time (ms) | Laser | Average laser illumination intensity |
|---|---|---|---|---|---|---|
| Figure 4 a-e<br>Supplementary Figure 16 & 17<br>Supplementary Video 6 & 7 | SST-Cre mice<br>AAV2/8-hSyn-Cre-WPRE-pA<br>Titer: 2.3 x $10^9$<br>AAV2/9-hSyn-FLEX-GCaMP6s-WPRE-pA<br>Titer: 5 x $10^{12}$ | 10 | 5 | 160 | 488 nm | 50 ~ 200 µm<br>0.73 $mW/mm^2$<br><br>150 ~ 300 µm<br>1.23 $mW/mm^2$<br><br>250 ~ 400 µm<br>1.23 $mW/mm^2$ |
| Figure 4f | SST-Cre mice<br>AAV2/8-hSyn-Cre-WPRE-pA<br>Titer: 5 x $10^9$<br>AAV2/9-hSyn-FLEX-GCaMP6s-WPRE-pA<br>Titer: 5 x $10^{12}$ | 10 | 5 | 160 | 488 nm | 50 ~ 200 µm<br>0.73 $mW/mm^2$<br><br>150 ~ 300 µm<br>1.23 $mW/mm^2$<br><br>250 ~ 400 µm<br>1.23 $mW/mm^2$ |
| Figure 5<br>Supplementary Video 8-10 | C57 | 8 | 70 | 4 | 635 nm | 0 ~ 150 µm<br>21.25 $mW/mm^2$<br><br>100 ~ 250 µm<br>25.3 $mW/mm^2$<br><br>200 ~ 350 µm<br>29 $mW/mm^2$<br><br>300 ~ 450 µm<br>32.8 $mW/mm^2$<br><br>400 ~ 550 µm<br>36.9 $mW/mm^2$ |

| | | | | | | |
|---|---|---|---|---|---|---|
| | | | | | | 500 ~ 650 μm<br>45.3 mW/$mm^2$ |
| Supplementary Figure 18 | SST-Cre mice<br>AAV2/8-hSyn-Cre-WPRE-pA<br>Titer: 2.3 x $10^9$<br>AAV2/9-hSyn-FLEX-GCaMP6s-WPRE-pA<br>Titer: 5 x $10^{12}$ | 10 | 5 | 160 | 488 nm | 0.73 mW/$mm^2$ |

**Supplementary Table 3| Acquisition parameters for two photon imaging in mice**

| Experiments | Mice strains/virus injection system | Age (weeks) | Frame rate (Hz) | Voxel size | Image size | Laser | Excitation laser power |
|---|---|---|---|---|---|---|---|
| Figure 4f | SST-Cre mice<br>AAV2/8-hSyn-Cre-WPRE-pA<br>Titer: $5 \times 10^9$<br>AAV2/9-hSyn-FLEX-GCaMP6s-WPRE-pA<br>Titer: $5 \times 10^{12}$ | 10 | 8 | 0.8 x 0.8 x 2.5 $\mu m^3$ | 1000 x 1000 | 920 nm | 0~250 μm<br>17.2 mW<br><br>250~400 μm<br>32.8 mW |

**Supplementary Video 1| Animated schematics of Confocal LFM with fast axial scanning**

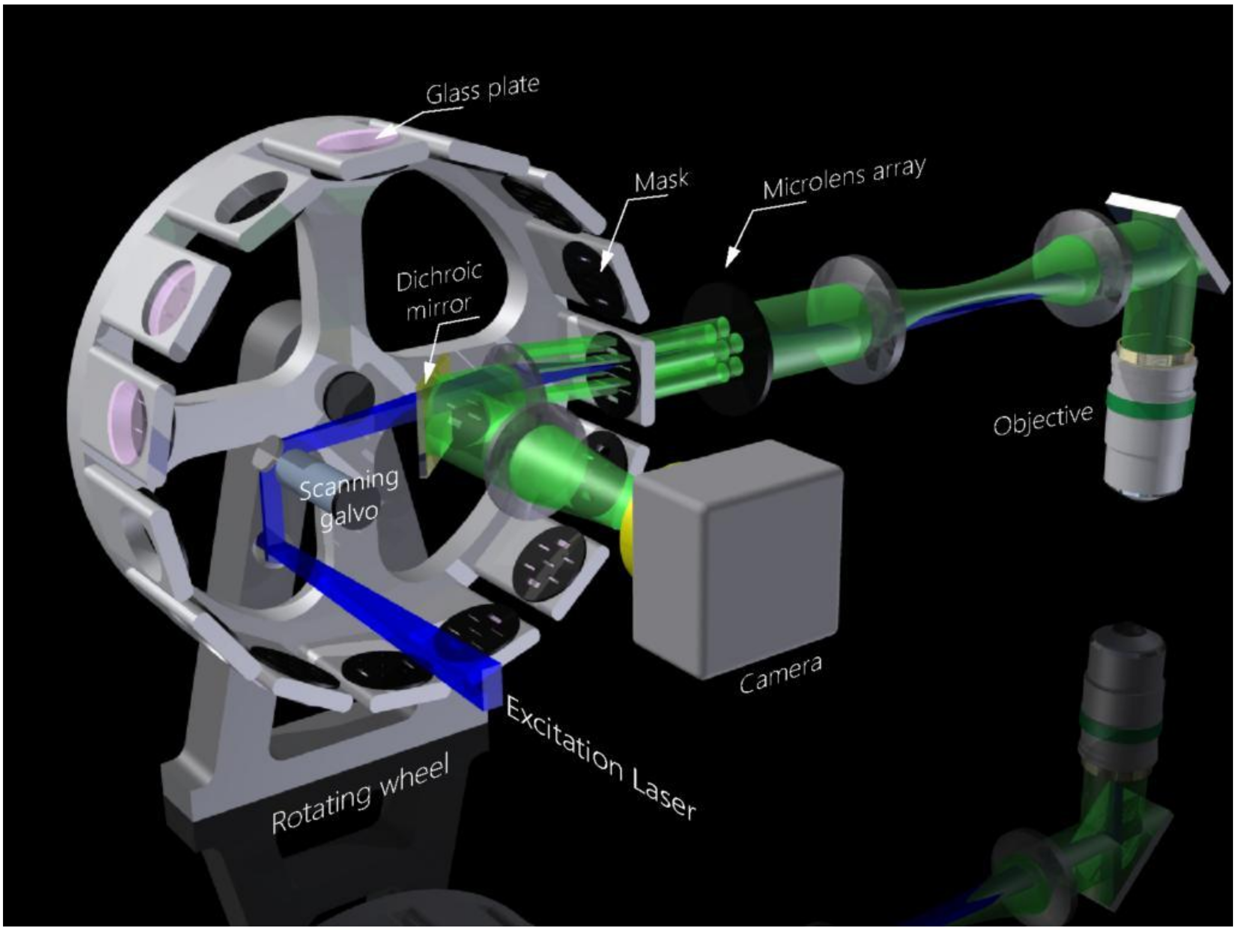


The excitation laser path and fluorescence imaging light path are in blue and green, respectively. The rotating wheel carrying masks and glass plates is rotated at constant speed. A trigger signal is generated by trigger slits on masks (Supplementary Figure 1 and 2, not shown here) to synchronize scanning galvo, laser and camera. Every time a trigger slit on a mask was rotated to the position where the trigger signal was generated, rotation of galvo mirror and rotating wheel was synchronized to make sure the excitation laser sheet went through a thin slit in the center of mask. Simultaneously, fluorescence signals collected by micro-lenses were spatially filtered by apertures on the same mask. More detailed descriptions of the system can be found in Figure 1 in the main text and methods.

**Supplementary Video 2| Comparison of Confocal LFM and non-confocal LFM when imaging spontaneous neural activities in restrained larval zebrafish brain**

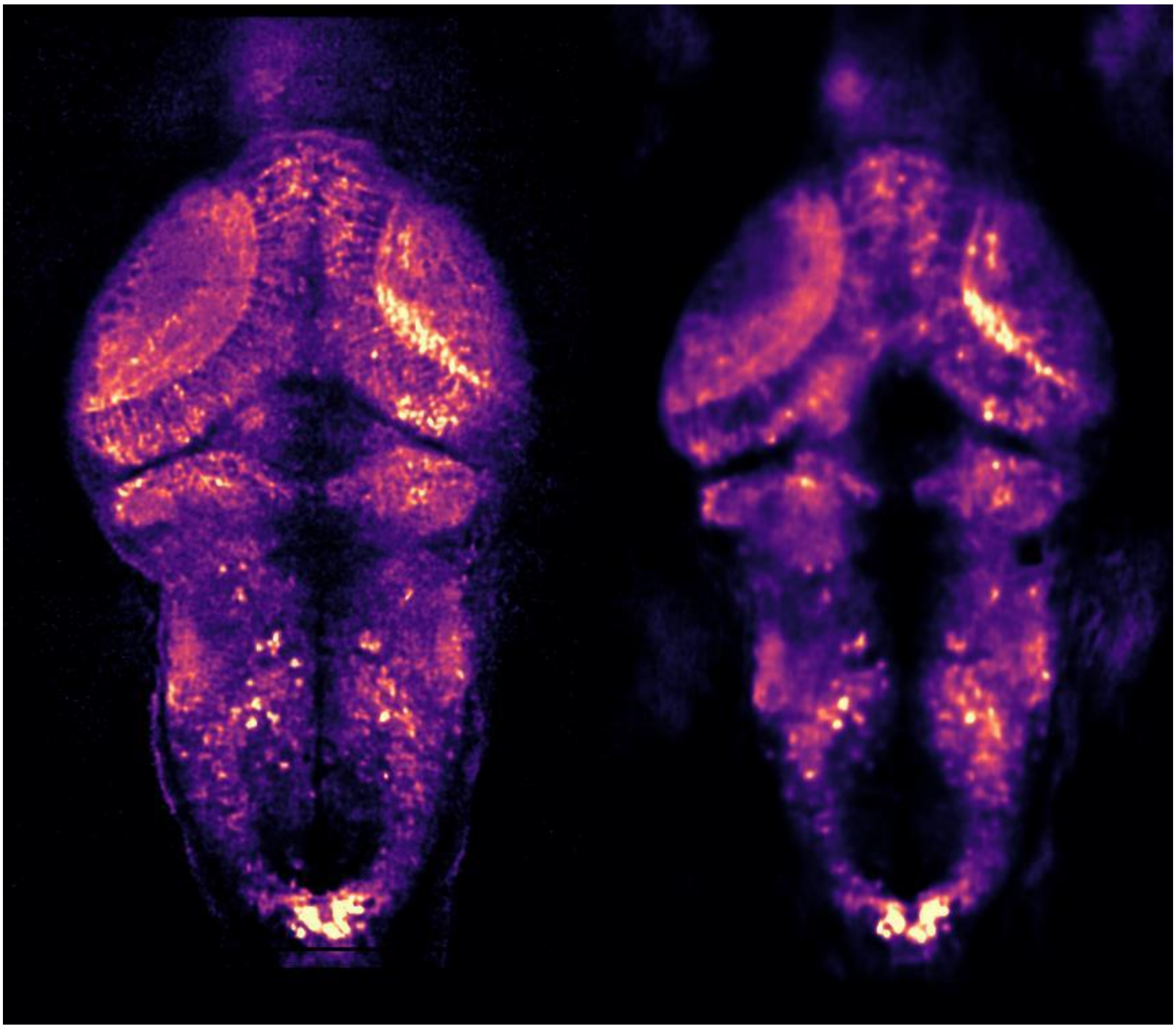

Spontaneous neural activities on a single plane in larval zebrafish brain (*Tg(HuC:GCaMP6s)*) imaged by Confocal LFM with (left) and without (right) confocal mask. Sampling rates were 1 Hz in both conditions. The larval zebrafish was 6 dpf with pan-neuronal labeling of GCamp6s localized in cytosol.

**Supplementary Video 3| 3D volumetric imaging of larval zebrafish brain**

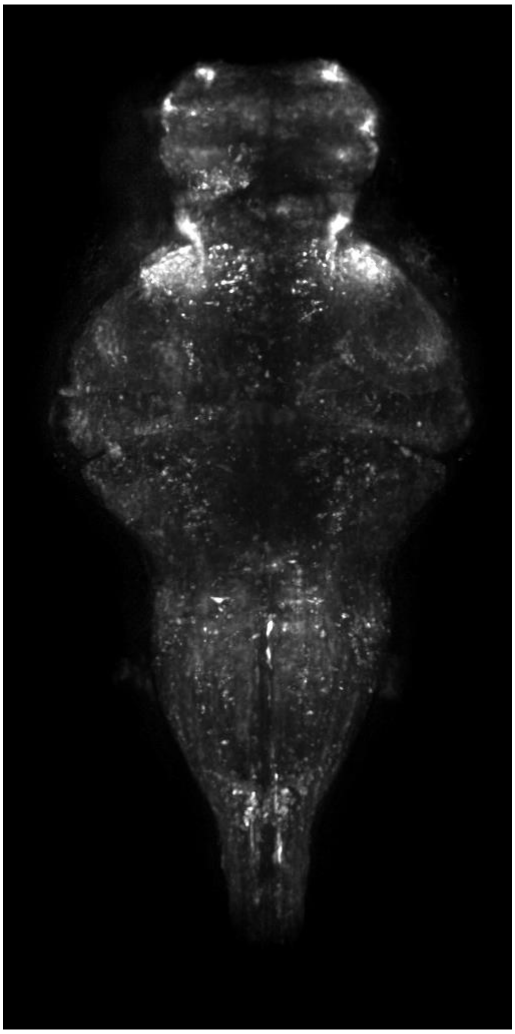

MIP in time of the calcium responses upon flash light stimuli in a larval zebrafish brain. The larval zebrafish is movement restrained. Five imaging volumes obtained sequentially from five different masks-glass plates combinations were stitched together to cover an entire imaging depth of ~200 μm. The same data was shown in Figure 2b, c & d in the main text and Supplementary Video 4.

**Supplementary Video 4| Whole brain functional imaging in larval zebrafish under light stimulation**

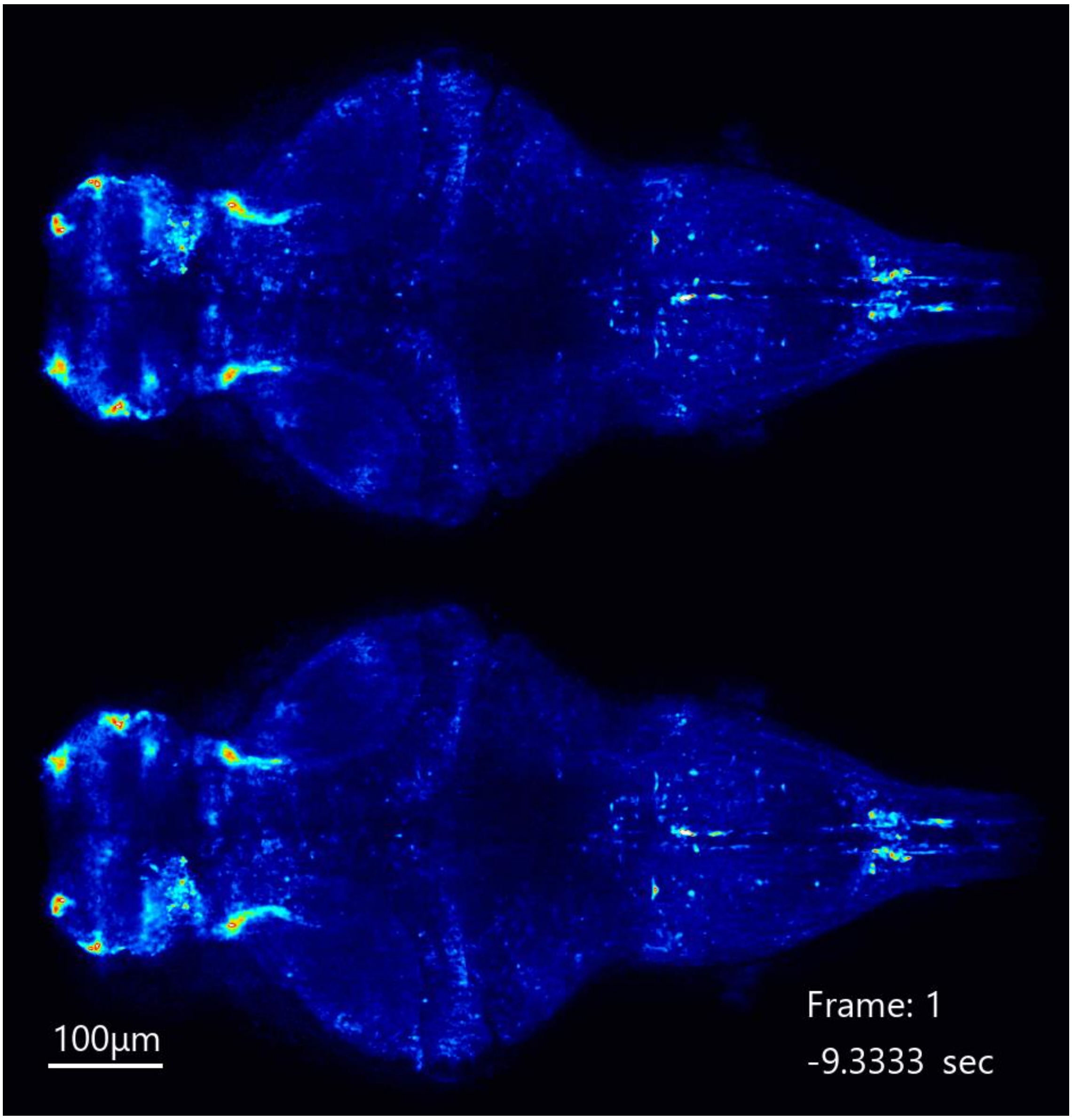


Larval zebrafish with pan neuronal labeling of cytosol-localized GCamp6s was imaged by Confocal LFM at 6 Hz volume rate. A flash of light was applied at time window of 0 — 4 s. The same data was shown in Figure 2b, c & d in the main text and Supplementary Video 3.

**Supplementary Video 5| Whole brain functional imaging of a freely swimming larval zebrafish during prey capture behavior**

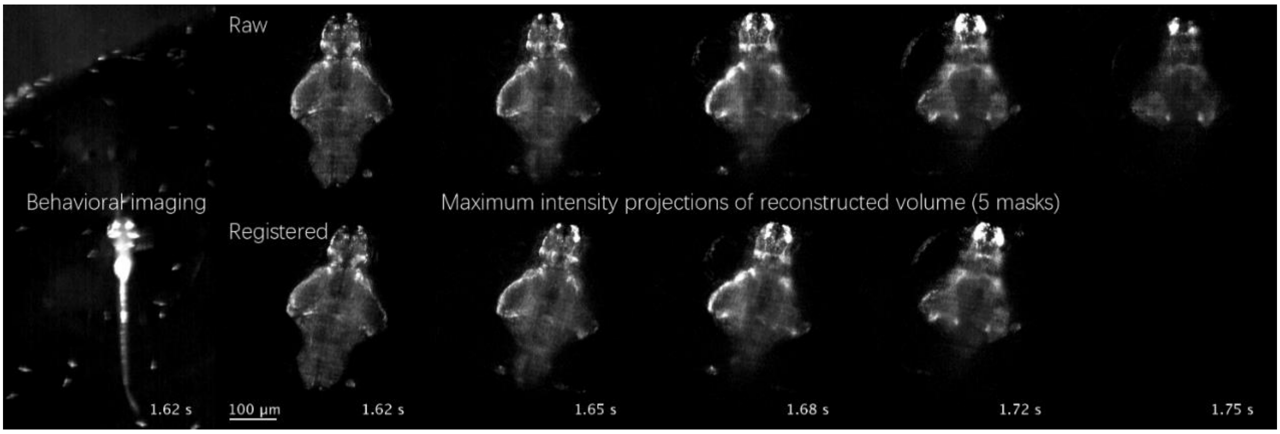


Confocal LFM imaging of a freely swimming larval zebrafish expressing cytosol-localized, pan-neuronal GCaMP6s (*Tg(HuC:GCaMP6*s), 8dpf). Left, behavioral recording of the zebrafish and paramecia. Right, MIPs of reconstructed volumes obtained in 5 different sets of masks and glass plates focusing at different depths. The raw reconstructions and registration of them were displayed on top and bottom, respectively. At 15.03 s, zebrafish larva initiated a hunting identified by its converging eyes onto the paramecium marked in blue and J-turn to align its heading direction. Its first try to capture the blue paramecium failed. At 15.63 s, it changed the target to the second paramecium marked in orange and captured it successfully. Video is playing in 4 times slower than real time. Scale bar, 100 μm. The same data was shown in Figure 3 in the main text.

**Supplementary Video 6| Unbiased volumetric reconstruction of active neural structures in awake mouse brain imaged by Confocal LFM**

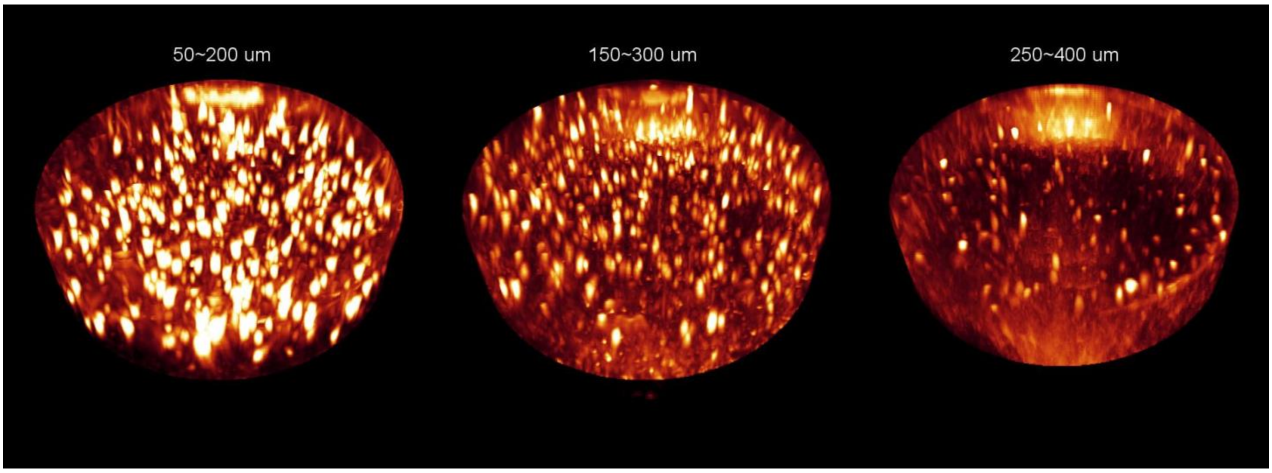


Volumetric rendering of spontaneously active neural structures imaged by Confocal LFM in awake mouse brain. Neurons in mouse brain were densely labeled with cytosol-localized GCamp6. The displayed images were standard deviations of unbiased volumetric reconstruction time series with 1600 time points. Detailed description of data processing can be found in Methods. Three volumes were captured at different depths in different sessions. The same data was shown in Figure 4a-e in the main text.

**Supplementary Video 7| Active neural structures extracted by CNMF-E on each z plane independently in an imaging volume captured by Confocal LFM**

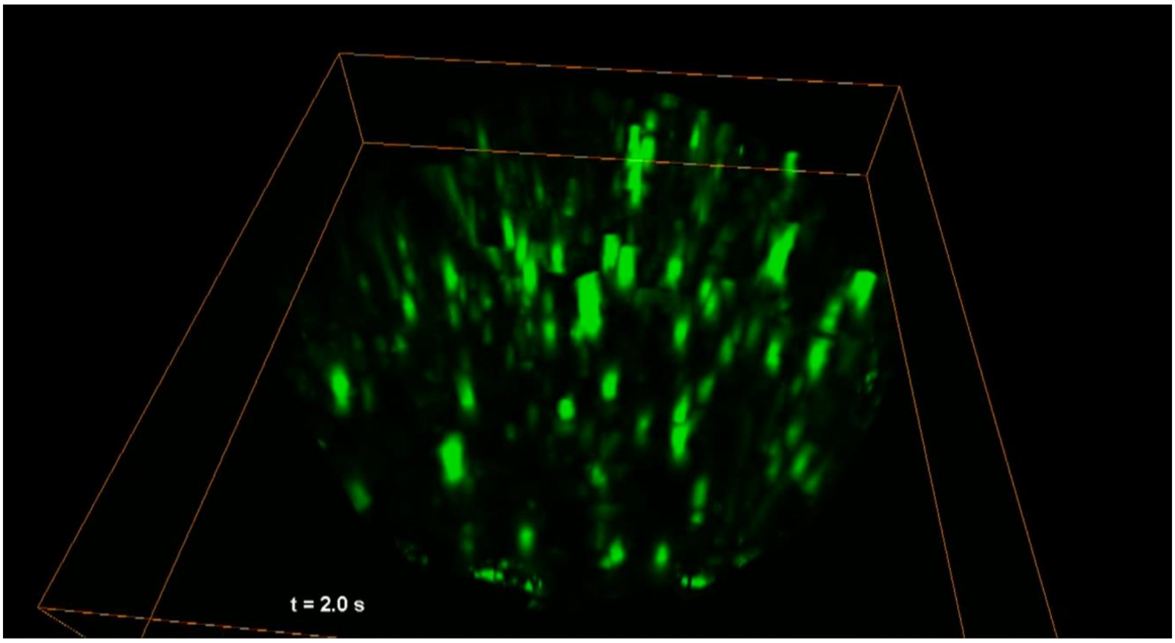


Volumetric rendering of time series of neural activities on their corresponding spatial footprints extracted by CNMF-E. The CNMF-E was separately applied on each z plane within the reconstructed volume (Methods). The imaging volume was from 100 μm ~ 200 μm below cortex surface. The spatial footprints of the same neuron at different z planes showed highly correlated temporal traces, which validated the CNMF-E extraction results. The same data was shown in Figure 4a-e in the main text.

**Supplementary Video 8| Imaging circulating blood cells in awake mouse brain by Confocal LFM**

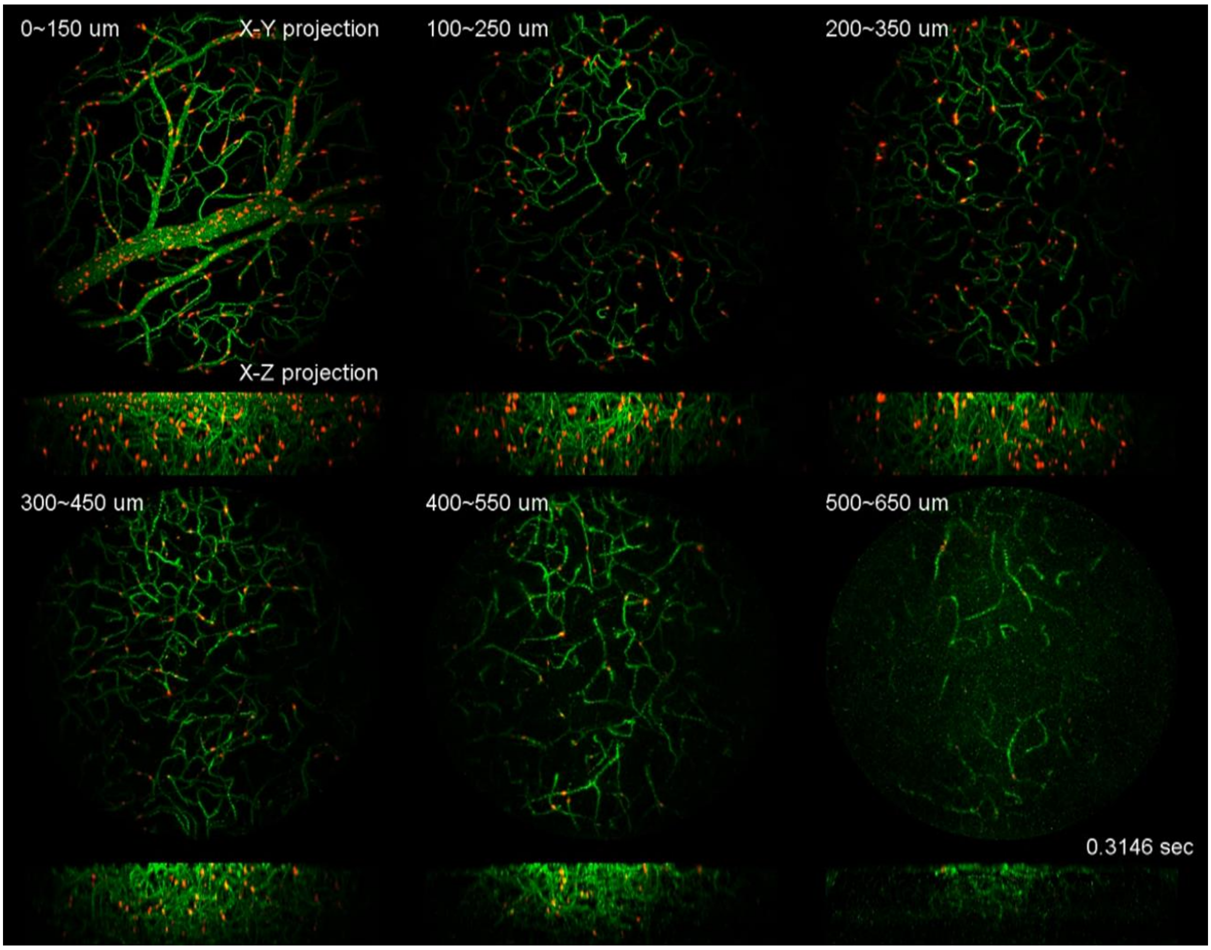


Imaging circulating blood cells in 6 volumes starting from surface to 650 μm at 70 Hz volume rate. Circulating cells labeled with red fluorescent dye were in red. Blood vessels, as found by averaging over time of images of labelled blood cells, were in green. Blood cells circulating as deep as 600 μm below the cortex surface could be imaged by Confocal LFM.

**Supplementary Video 9| Imaging circulating blood cells in awake mouse brain over extended period of time**

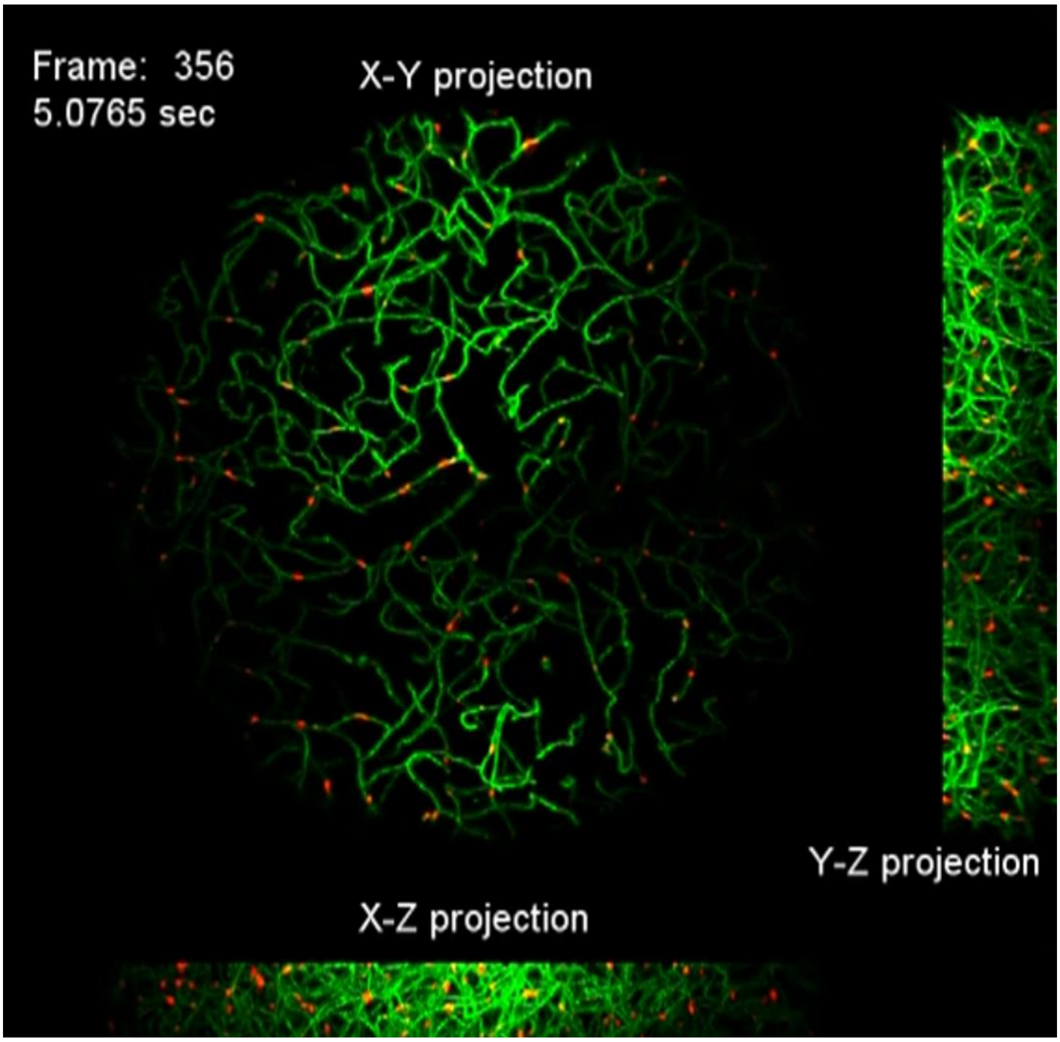


Volumetric imaging of circulating blood cells 100 µm ~ 250 µm below the mouse cortex surface over entire imaging session (1260 frames in 18 s). This data is the same as the second volume in Supplementary Video 8 but with more time points.

**Supplementary Video 10| Analysis of instantaneous speeds of circulating cells in blood vessels**

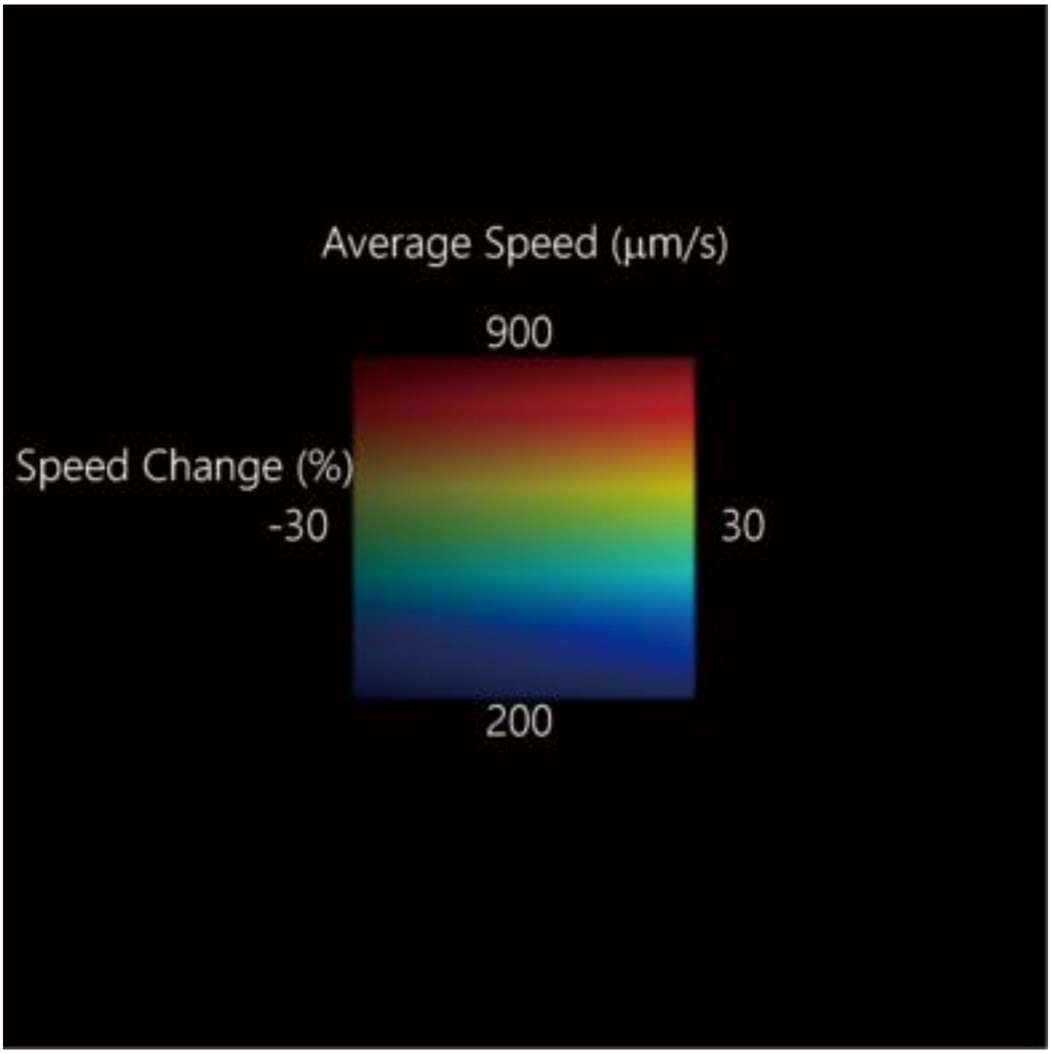


By tracking single cells in blood vessels, their speeds can be inferred. The skeletons of blood vessels were colored based on the average speeds of circulating blood cells in them over the entire imaging session (1260 frames in 18 s). The change of brightness on each blood vessel skeleton indicated the change of instantaneous speeds of cells in it. This analysis was carried on the imaging volume covering 100 μm ~ 250 μm below mouse cortex surface ( same as in Supplementary Video 9). The skeletons of blood vessels in the volume were projected in z direction for display.
